# Independent domestication events in the blue-cheese fungus *Penicillium roqueforti*

**DOI:** 10.1101/451773

**Authors:** Emilie Dumas, Alice Feurtey, Ricardo C Rodríguez de la Vega, Stéphanie Le Prieur, Alodie Snirc, Monika Coton, Anne Thierry, Emmanuel Coton, Mélanie Le Piver, Daniel Roueyre, Jeanne Ropars, Antoine Branca, Tatiana Giraud

## Abstract

Domestication provides an excellent framework for studying adaptive divergence. Using population genomics and phenotypic assays, we reconstructed the domestication history of the blue cheese mold *Penicillium roqueforti.* We showed that this fungus was domesticated twice independently. The population used in Roquefort originated from an old domestication event associated with weak bottlenecks and exhibited traits beneficial for pre-industrial cheese production (slower growth in cheese and greater spore production on bread, the traditional multiplication medium). The other cheese population originated more recently from the selection of a single clonal lineage, was associated to all types of blue cheese worldwide but Roquefort, and displayed phenotypes more suited for industrial cheese production (high lipolytic activity, efficient cheese cavity colonization ability and salt tolerance). We detected genomic regions affected by recent positive selection and putative horizontal gene transfers. This study sheds light on the processes of rapid adaptation and raises questions about genetic resource conservation.

## Introduction

What are the mechanisms of adaptive divergence (population differentiation under selection) is a key question in evolutionary biology for understanding how organisms adapt to their environment and how biodiversity arises. Domestication is a special case of adaptive divergence, involving strong and recent selection for traits that can be easily identified. Furthermore, closely related non-domesticated populations are often available, making it possible to contrast their traits and genomes with those of domesticated populations. Studying domestication can therefore provide a deeper understanding of the mechanisms of adaptive divergence. This approach has proved to be powerful for reconstructing the history of divergence and the genetic architecture of traits selected by humans when applied to maize and teosinte or to dog breeds and wolves (Albert et al., 2012; Axelsson et al., 2013; Freedman et al., 2016; Hake and Ross-Ibarra, 2015; Li et al., 2016; Wang et al., 2015) Comparisons of domesticated varieties selected for different phenotypes have also proved to be a powerful approach for elucidating the mechanisms of adaptation, for example in dog breeds and pigeons (Parker et al., 2017; Shapiro et al., 2013)]. Studies on genetic diversity and subdivision in domesticated organisms provides also crucial information for the conservation of genetic resources. Indeed, recent breeding programs have resulted in a massive loss of genetic diversity in crops and breeds, potentially jeopardizing adaptive potential for improvement (Gouyon et al., 2010; Harlan, 1992; Vavilov, 1992).

Fungi are interesting eukaryotic models for adaptive divergence studies, with their small genomes, easy access to the haploid phase and experimental tractability for *in vitro* experiments (Giraud et al., 2017; Gladieux et al., 2014). Many fungi are used as food sources (Dupont et al., 2016) and some have been domesticated for food production. Propagation of the latter is controlled by humans, and this has resulted in genetic differentiation from wild populations (Almeida et al., 2017, 2014; Gallone et al., 2016; Gibbons et al., 2012; Gonçalves et al., 2016) and the evolution of specific phenotypes beneficial for humans (Dupont et al., 2016; Gallone et al., 2016; Gibbons et al., 2012; Gibbons and Rinker, 2015; Marsit et al., 2015). *Saccharomyces cerevisiae* yeasts domesticated for fermentation have provided important insight into adaptive divergence mechanisms, with different yeast lineages independently domesticated for different usages (Borneman et al., 2011; Gonçalves et al., 2016; Peter et al., 2018). Studies about yeast adaptation for alcohol and cheese production have highlighted the proximal genomic mechanisms involved, including horizontal gene transfer, selective sweep, hybridization and introgression (Legras et al., 2018; Marsit et al., 2015; Morales and Dujon, 2012; Novo et al., 2009; Peter et al., 2018).

*Penicillium roqueforti,* a filamentous fungus used in the dairy industry to impart the typical veins and flavor of blue cheeses, has recently emerged as an excellent model for studying adaptive divergence (Cheeseman et al., 2014; Ropars et al., 2015). Blue cheeses, including Roquefort, Gorgonzola and Stilton, are highly emblematic foods that have been produced for centuries (Vabre, 2015). The strongest genetic subdivision reported in *P. roqueforti* concerns the differentiation of a cheese-specific population that has acquired faster growth in cheese than other populations and better excludes competitors, thanks to very recent horizontal gene transfers, at the expense of slower growth on minimal medium (Gillot et al., 2015; Ropars et al., 2015, 2017). Such genetic differentiation and recent acquisition of traits beneficial to cheesemaking in *P. roqueforti* suggests genuine domestication, i.e., adaptation under selection by humans for traits beneficial for food production. A second population identified in *P. roqueforti* and lacking the horizontally-transferred regions includes strains isolated from cheese and other environments, such as silage, lumber and spoiled food (Gillot et al., 2015; Ropars et al., 2014, 2017). *Penicillium roqueforti* is the main contaminant of silage, spoilage typically occurring following breaks in plastic or after opening the stack for cattle feeding. In this context, it can produce harmful mycotoxins causing health disorder in cattle (Malekinejad et al., 2015). In addition, *P. roqueforti* is one of the most common *Penicillium* species in spoiled food, where it is also responsible for mycotoxin production (Rundberget et al., 2004). The existence of further genetic subdivision separating populations according to the original environment, or protected designation of origin (PDO) for cheese strains has been suggested, but, because it was based only on a few microsatellite markers, the resolution power was low (Gillot et al., 2015; Ropars et al., 2014, 2017). Secondary metabolite production (aroma compounds and mycotoxins) and proteolysis activity have been shown to differ between strains from different PDOs (Gillot et al., 2017). A high-quality *P. roqueforti* genome reference is available (Cheeseman et al., 2014), allowing more powerful analyses based on population genomics.

Another asset of *P. roqueforti* as an evolutionary model is the availability of vast collections of cheese strains and of historical records concerning cheesemaking (Aussibal, 1983; Labbe and Serres, 2009, 2004; Marre, 1906; Marres, 1935; Vabre, 2010). While the presence of *P. roqueforti* in cheeses was initially fortuitous, since the end of the 19^th^ century, milk or curd has been inoculated with the spores of this fungus for Roquefort cheese production. Spores were initially multiplied on bread, before the advent of more controlled *in vitro* culture techniques in the 20^th^ century (Aussibal, 1983; Labbe and Serres, 2009, 2004; Marre, 1906; Marres, 1935; Vabre, 2010). Bread was inoculated by recycling spores from the best cheeses from the previous production (i.e., back-slopping) (Aussibal, 1983; Labbe and Serres, 2009, 2004; Marre, 1906; Marres, 1935; Vabre, 2010). This corresponds to yearly selection events since the 19^th^ century until ca. 20 years ago when strains were stored in freezers. After World War II, strains were isolated in the laboratory for industrial use and selected based on their technological and organoleptic impact in cheeses and compounds produced (Besana et al., 2017), which have likely accelerated domestication. This history further suggests that there may have been genuine domestication, i.e., an adaptive evolution triggered by human selection for cheese quality. Unintentional selection may also have been exerted on other traits, including growth and spore production on bread, the traditional multiplication substrate.

By sequencing multiple *P. roqueforti* genomes from different environments and analyzing large collections of cheese strains, we provide evidence for adaptive divergence. We identified four genetically differentiated populations, two including only cheese strains and two other populations including silage- and food-spoiling strains. We inferred that the two cheese populations corresponded to two independent domestication events. The first cheese population corresponded to strains used for Roquefort production and arose through a weaker and older domestication event, with multiple strains probably originating from different cultures on local farms in the PDO area, presumably initially selected for slow growth before the invention of refrigeration systems. The second cheese population experienced an independent and more recent domestication event associated with a stronger genetic bottleneck. The non-Roquefort cheese population showed beneficial traits for modern industrial production of cheese (e.g. faster growth in salted cheese, more efficient cheese cavity colonization and faster lipid degradation activities), while the Roquefort cheese population showed greater spore production on bread, the traditional medium for spore production. The four populations further showed differences in proteolysis activities, with a higher variance in the cheese populations. The two cheese populations also had different volatile compound profiles, with likely effects on cheese flavor. These phenotypic differences might be associated with genomic regions affected by recent positive selection and genomic islands specific to a single cheese population. Some of these genomic regions may have been acquired by horizontal gene transfers and have putative functions in the biochemical pathways leading to the development of cheese flavor.

## Results

### Two out of four populations are used for cheesemaking: one specific to the Roquefort PDO and a worldwide clonal population

We sequenced the genomes of 34 *P. roqueforti* strains from public collections (Ropars et al., 2017), including 17 isolated from blue cheeses (e.g., Roquefort, Gorgonzola, Stilton), 17 isolated from non-cheese environments (mainly spoiled food, silage, and lumber), and 11 outgroup genomes from three *Penicillium* species closely related to *P. roqueforti* (Supplementary Table 1). After data filtering, we identified a total of 115,544 SNPs from the reads mapped against the reference *P. roqueforti* FM164 genome (29×10^6^ bp, 48 scaffolds).

We used three clustering methods free from assumptions about mating system and mode of reproduction, based on genetic differences (principal component analyses, SplitsTree and clustering based on similarities between genotypes along the genomes in 50 SNP-windows). The three methods separated the *P. roqueforti* strains into four genetic clusters (Figs. 1, 2 and 3), two of which almost exclusively contained cheese strains (the exceptions being two strains isolated from a brewery and brioche, Figs. 1 and 2, probably corresponding to feral strains). One cluster contained both silage strains (N=4) and food-spoiling strains (N=4), and the last cluster contained mostly food-spoiling strains (N=5) plus strains from lumber (N=2) (Figs. 1 and 2, and Supplementary Table 1). Noteworthy, these two clusters corresponding to strains from other environments did not include a single cheese strain. The two cheese clusters were not the most closely related one to each other, suggesting independent domestication events (Figs. 1 and 2). Moreover, cheese clusters displayed much lower genetic diversity than non-cheese clusters, as shown by their small ϴ values (corresponding to 4N_e_µ, i.e., the product of the effective population size and the mutation rate) and more homogeneous colors in distance-based clustering (Table 1 and Fig. 2). One of the two cheese clusters displayed a particularly low level of genetic diversity (Table 1 and Fig. 2) with only 0.03% polymorphic sites, and a lack of recombination footprints (i.e., a higher level of linkage disequilibrium, as shown by the more gradual decay of r² values (Supplementary Fig. 1), and by the large single-color blocks along the genomes, Fig. 2). These findings suggest that the second cheese population is a single clonal lineage. The first cheese population also appears to lack recombination footprints, while including several clonal lineages (Fig. 2). Given such a lack of recombination footprints, clustering methods free of assumptions on modes of recombination were better suited to analyse the dataset. The Structure software, that assumes random mating, nevertheless yielded similar results (Supplementary Fig. 2).

**Figure 1:**
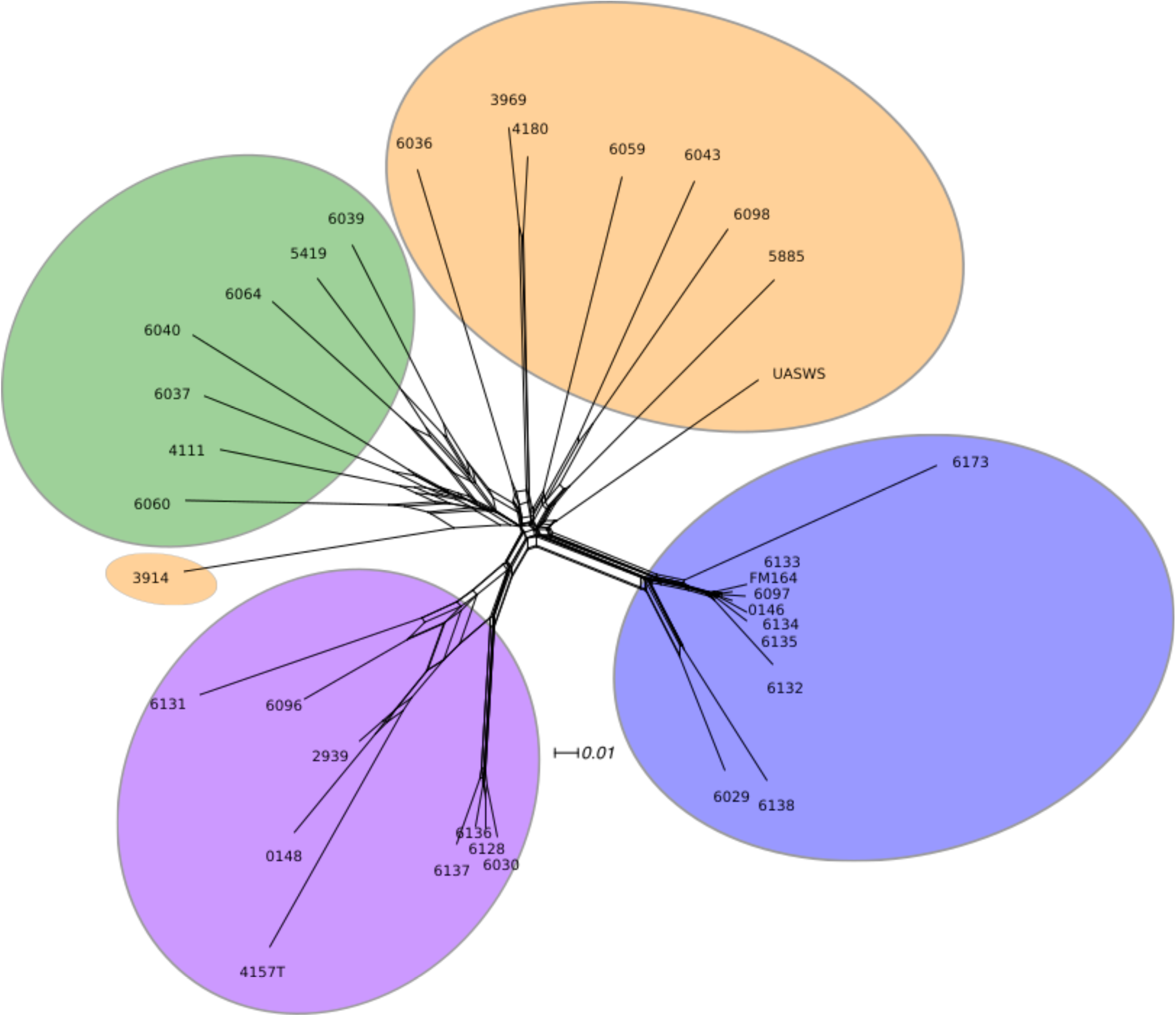
Diversity and population subdivision in *Penicillium roqueforti*. Unrooted phylogenetic network of P. roqueforti strains generated with SplitsTree4 from SNP variation. The scale bar indicates the number of substitutions per site. The letters indicate the origin of the strains, C = cheese, F = spoiled food, S = silage and L = lumber. The color indicates assignment to one of the four *P. roqueforti* populations identified, as in the other figures. Blue, non-Roquefort; purple, Roquefort; green, lumber/food spoilage, and; orange, silage/food spoilage.

**Figure 2:**
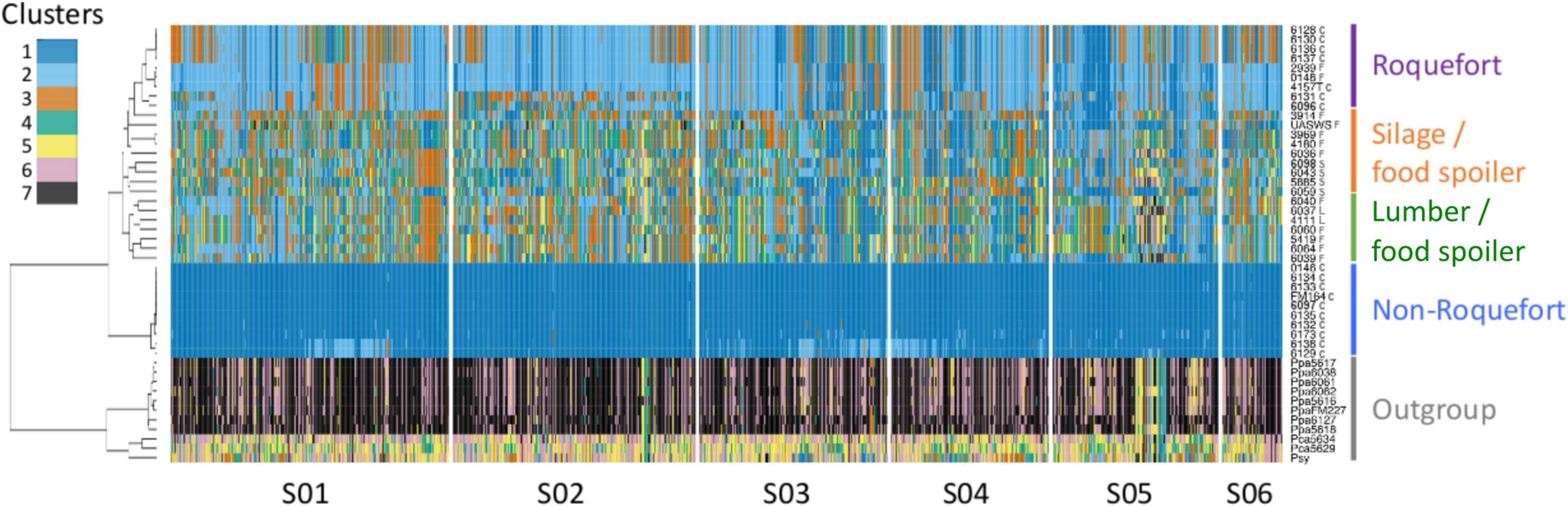
Clustering of *Penicillium roqueforti* along the FM164 reference genome using non-overlapping 50 SNP sliding windows. Clustering was done in each window using the “mclust” function with Gaussian mixture modelling and using the Gower’s distance between haplotypes. The maximum number of clusters was fixed to seven, corresponding to the three outgroup species plus the four populations of *P. roqueforti*. Each color corresponds to a cluster. Windows containing fewer than 50 SNPs at the edge of scaffolds are not represented. The dendrogram on the left side was reconstructed using hierarchical clustering based on the Gower’s distance between clusters for the entire genome. The histogram on the top left represents the distribution of the number of clusters inferred for the whole genome. The letters indicate the origin of the strains, C = Cheese, F = Food, S = Silage and L = Lumber.

**Figure 3:**
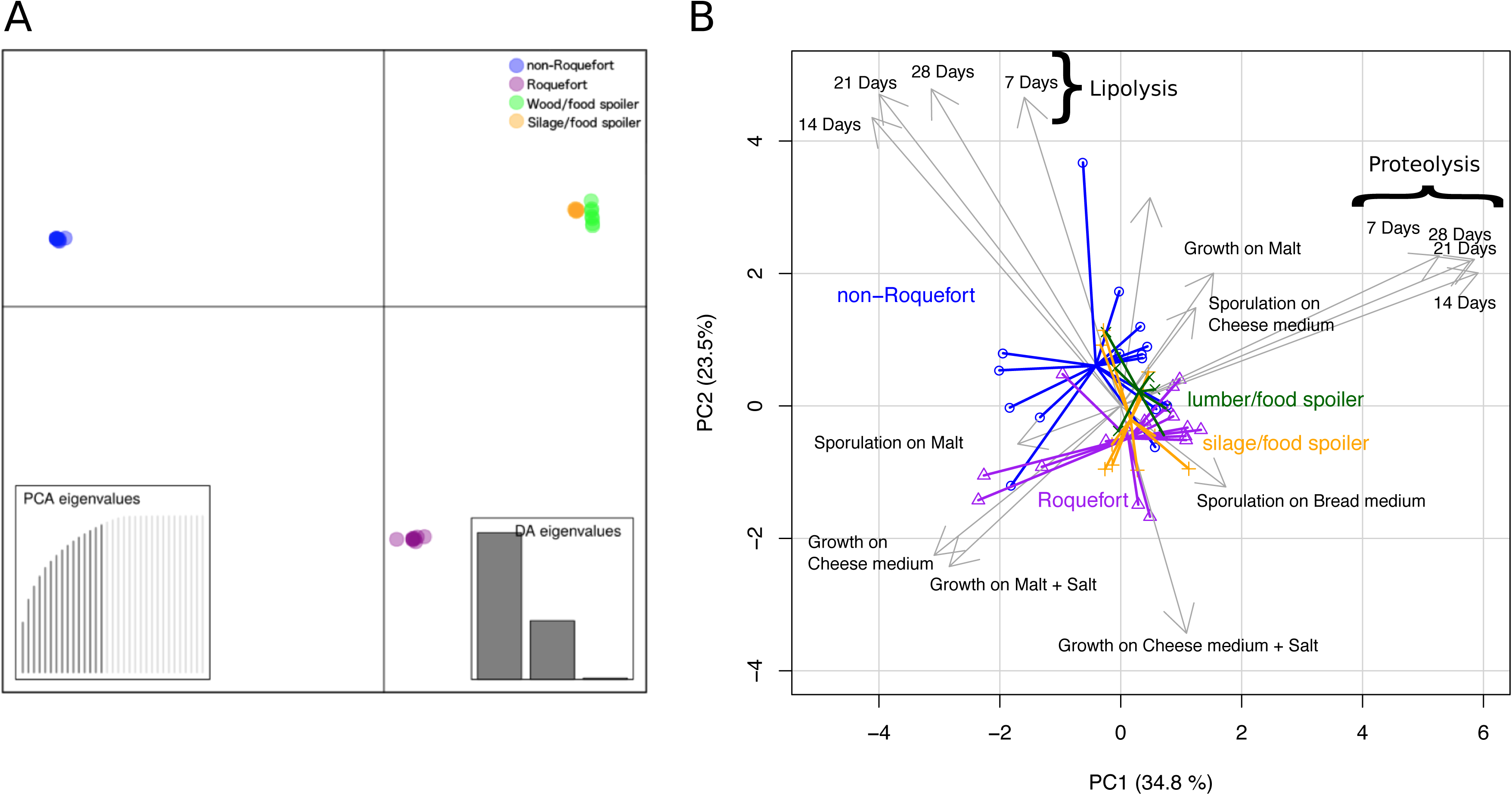
Genetic and phenotypic differentiation among *Penicillium roqueforti* populations. Colors correspond to the genetic clusters as in other figures. A: genetic differentiation assessed by a discriminant analysis of principal components (DAPC) based on genome-wide single-nucleotide polymorphisms (SNPs). The dots represent the strains and the colors the four populations identified based on the genealogical tree in Fig. 1 as well as the similarity clustering in Fig. 2. The insets show the distribution of eigenvalues for the principal component analysis (PCA) and for the discriminant analysis (DA). B: phenotypic differentiation among *P. roqueforti* genetic clusters illustrated by a PCA based on all tested phenotypes. Colors correspond to the genetic clusters as in other figures. Missing data correction has been done using Bayesian correction in the pcaMethods package (Stacklies et al., 2007).

**Table 1:**
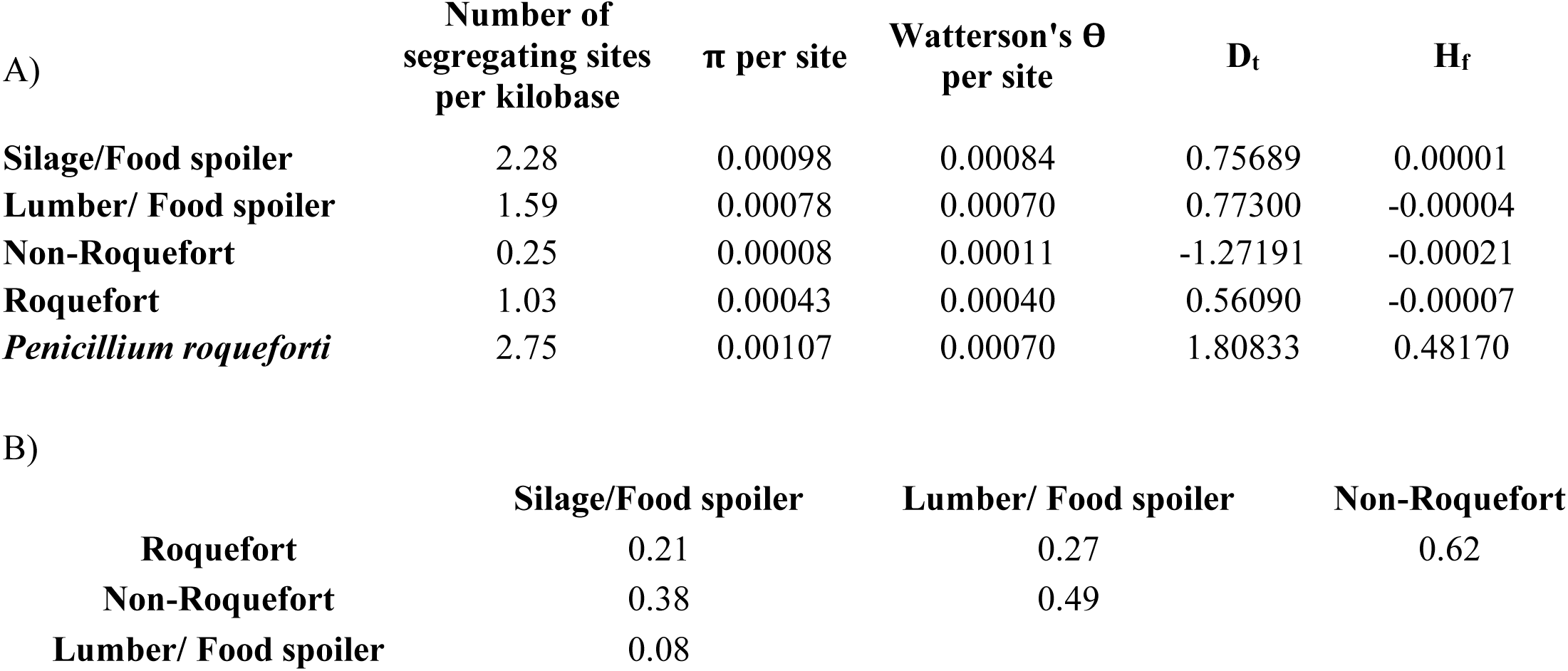
Population genetics statistics in the four Penicillium roqueforti populations. A: Statistics calculated by averaging values on 1144 sliding windows of 50 kb with 25 kb overlap. B: F_ST_ values calculated on pairwise comparisons.

We used genome sequences to design genetic markers (Supplementary Table 2) for assigning a collection of 65 strains provided by the main French supplier of *P. roqueforti* spores for artisanal and industrial cheesemakers, 18 additional strains from the National History Museum collection in Paris (LCP) and 31 strains from the collection of the Université de Bretagne Occidentale (UBOCC, Supplementary Table 1) to the four genetic clusters. Out of these 148 strains, 55 were assigned to the more genetically diverse of the two cheese clusters. The majority of these strains included strains used for Roquefort PDO cheese production (N=30); three strains originated from Bleu des Causses cheeses (Supplementary Fig. 3, Supplementary Table 1), produced in the same area as Roquefort and using similarly long storage in caves. The remaining strains of this cluster included samples from other blue cheeses (N=13), unknown blue cheeses (N=5) or other environments (N=4), the latter likely associated with feral strains. Because of the strong bias of usage toward Roquefort production, we refer to this cluster hereafter as the “Roquefort population”. Of the remaining 95 strains, 60 belonged to the second cheese cluster, which was less genetically diverse and contained mainly commercial strains used to produce a wide range of blue cheeses (Supplementary Fig. 3, Supplementary Table 1). This cluster was therefore named the “non-Roquefort population”. A single strain (LCP00146) in this non-Roquefort population had been likely sampled from a Roquefort cheese, but it did not appear phenotypically different from other strains in its genetic group; the “Roquefort” origin may however be dubious as no brand was recorded for this strain from an old collection. The Roquefort population also included 13 strains used to inoculate other types of blue cheese (e.g. Gorgonzola or Bleu d’Auvergne), but strains from these types of cheeses were more common in the non-Roquefort population. The non-Roquefort cluster contained strains harbouring *Wallaby* and *CheesyTer*, two large genomic regions recently shown to have been transferred horizontally between different *Penicillium* species from the cheese environment and conferring faster growth on cheese (Cheeseman et al., 2014; Ropars et al., 2015), whereas all the strains in the Roquefort cluster lacked those regions.

### Two independent domestication events in Penicillium roqueforti for cheesemaking

We compared 11 demographic scenarios with approximate Bayesian computation (ABC), simulating either a single domestication event (the most recent divergence event then separating the two cheese populations) or two independent domestication events, with different population tree topologies and with or without gene flow (Supplementary Fig. 4). Parameters in the scenarios modeled corresponded to the divergence dates, the strength and dates of bottlenecks and population growth, and rates of gene flow. ABC simulates sequence evolution under the various scenarios using the coalescent theory framework and compares various population statistics under a Bayesian framework between the simulation outputs and the observed data to identify the most likely scenario (Beaumont et al., 2002). The ABC results showed that the two *P. roqueforti* cheese populations (Roquefort and non-Roquefort) resulted from two independent domestication events (Fig. 4). The highest posterior probabilities were obtained for the S4 scenario, in which the two cheese populations formed two lineages independently derived from the common ancestral population of all *P. roqueforti* strains (Fig. 4, model choice and parameter estimates in Supplementary Fig. 4). We inferred much stronger bottlenecks in the two cheese populations than in the non-cheese populations, with the most severe bottleneck found in the non-Roquefort cheese population. Some gene flow (m=0.1) was inferred between the two non-cheese populations but none with cheese populations. The bottleneck date estimates in ABC had too large credibility intervals to allow inferring domestication dates (Supplementary Fig. 4E). We therefore used the multiple sequentially Markovian coalescent (MSMC) method to estimate times since domestication, considering that they corresponded to the last time there was gene flow between genotypes within populations, given the lack of recombination footprints in cheese population and the mode of conservation and clonal growth of cheese strains by humans, and given that this also corresponds to bottleneck date estimates in coalescence. The domestication for the Roquefort cheese population was inferred seven times longer ago than for the non-Roquefort cheese population, both domestication events being recent (ca. 760 versus 140 generations ago, Fig. 5B-C). Unfortunately, generation time, and even generation definition, are too uncertain in the clonal *P. roqueforti* populations to infer domestication dates in years. In addition, the MSMC analysis detected two bottlenecks in the history of the Roquefort cheese population (Fig. 4B).

**Figure 4:**
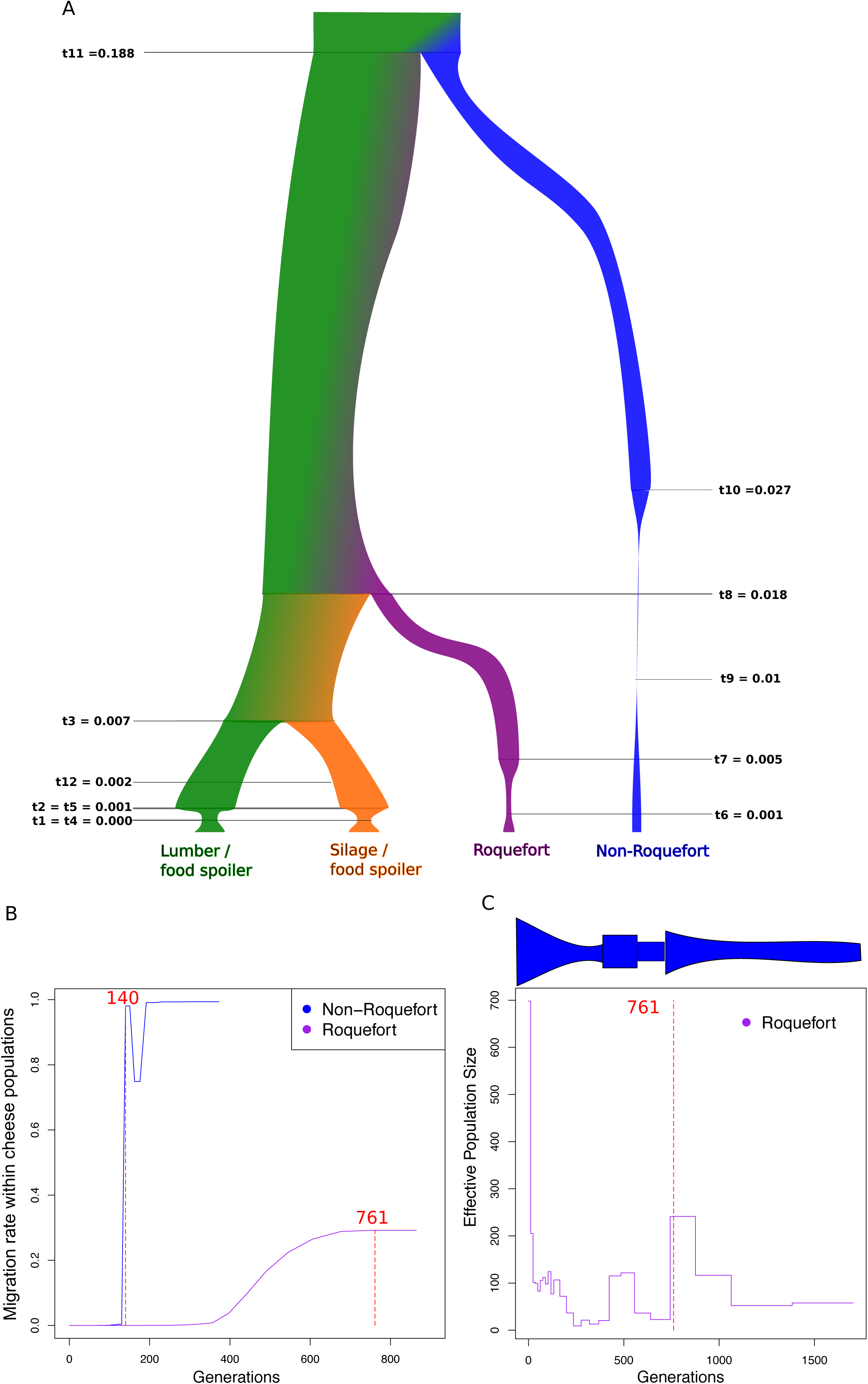
Demographic history of *Penicillium roqueforti* populations. A. Demographic scenario (S4) with the highest posterior probability for the history of *Penicillium roqueforti* populations. Estimates of time since divergence are indicated in units of 2N_e_ generations (Supplementary Figure 4 E); effective population sizes and their variation (bottlenecks) are represented by the widths of the genealogy branches, with relative sizes being represented to scale. The color indicates assignment to the *P. roqueforti* populations as in the other figures. B. Estimated past migration rate (gene flow) within each of the two cheese populations backward in time (t=0 represents the present time). The dashed red lines represent the inferred times of domestication, estimated as the last time gene flow occurred within cheese populations. C. Estimated demographic history for the Roquefort population using the multiple sequentially Markovian coalescent (MSMC) method. The inferred population effective size is plotted along generations backward in time (t=0 represents the present time). The dashed red line represents the inferred domestication time, estimated as the last time gene flow occurred within the Roquefort population (Fig. 4B). The scheme above the figure represents a schematic view of the effective population size along generations, representing the two bottlenecks.

**Figure 5:**
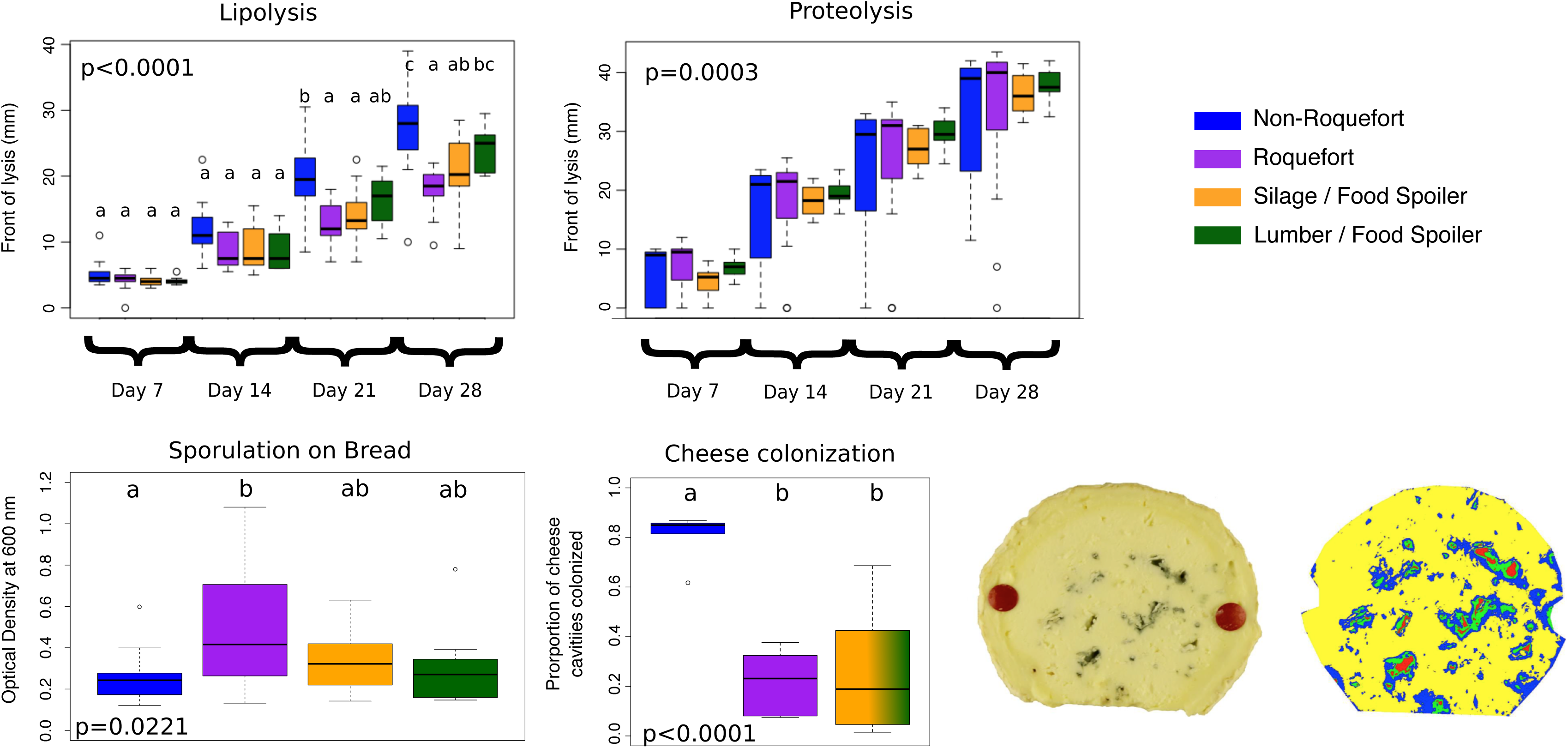
Differences in phenotype between *Penicillium roqueforti* populations for various traits relevant for cheesemaking. The color indicates assignment to the *P. roqueforti* populations identified, as in the other figures. Horizontal lines on the boxplots represent the upper quartile, the median and the lower quartile. Dots represent the outlier values. Different letters indicate significant differences (Supplementary Table 4). **A**: Lipolytic activity measured at four different dates; **B**: Proteolytic activity measured at four different dates; **C**: Spore production on bread medium measured as optical density by spectrophotometer; **D**: Cheese cavity occupation (i.e., percentage of total cheese cavity space colonized by the fungus, as measured on images) estimated in experimental cheeses by image analysis. The two clusters of non-cheese strains were pooled, as there were too few strains per cluster to test differences between the lumber/food spoiler and silage/food spoiler clusters. (a) Picture of a cheese slice. (b) Corresponding image analysis using the geospatial image processing software ENVI (Harris Geospatial Solution). Colors correspond to pixel classification based on their color on the picture. In yellow and blue: the inner white part of the cheese; in green and red: cavities.

### Contrasting fitness traits between cheese populations

We tested whether different phenotypes relevant for cheesemaking had evolved in the two cheese clusters, relative to other populations (Fig. 5, Supplementary Table 3). We first produced experimental cheeses inoculated with strains from the different *P. roqueforti* populations to assess their ability to colonize cheese cavities, a trait that may have been subject to human selection to choose inocula producing the most visually attractive blue cheeses. The fungus requires oxygen and can therefore sporulate only in the cheese cavities, its spores being responsible for the typical color of blue-veined cheeses; the application of highly salted solutions followed by tin foil wrapping prevents sporulation on the surface of cheeses. Strains from the non-Roquefort cheese population were the most efficient colonizers of cheese cavities (Supplementary Table 4); no difference was detected between strains from the Roquefort and non-cheese populations (Fig. 5).

As *P. roqueforti* strains were traditionally multiplied on bread loaves for cheese inoculation, they may have been subject to unintentional selection for faster growth on bread. However, growth rate on bread did not significantly differ between populations (Fig. 5, Supplementary Table 4).

We then assessed lipolytic and proteolytic activities in the *P. roqueforti* populations. These activities are important for energy and nutrient uptake, as well as for cheese texture and the production of volatile compounds responsible for cheese flavors (Gillot et al., 2017; McSweeney, 2004). Lipolysis was faster in the non-Roquefort cheese population than in the Roquefort and silage/food spoiling populations (Fig. 5, Supplementary Table 4). A strong population effect was found for proteolytic activity (Supplementary Table 4), with faster proteolysis activities in cheese populations (Fig. 5), although posthoc pairwise tests were not significant. Variances showed significant differences between populations (Levene test F-ratio=5.97, d.f.=3, P<0.0017), with the two cheese populations showing the highest variances, and with extreme values above and below those in non-cheese populations (Fig. 5). Noteworthy, proteolysis is a choice criterion for making different kinds of blue cheeses that is often showcased by culture producers (e.g. https://www.lip-sas.fr/index.php/nos-produits/penicillium-roquefortii/18-penicillium-roquefortii). This suggests that some cheese strains may have been selected for higher and others for lower proteolytic activity. Alternatively, selection could have been relaxed on this trait in the cheese populations, leading to some mutations decreasing and other increasing proteolysis in different strains, thus increasing variance in the populations.

The ability of *P. roqueforti* strains to produce spores may also have been selected by humans, both unwittingly, due to the collection of spores from moldy bread, and deliberately, through the choice of inocula producing bluer cheeses. We detected no difference in spore production between the *P. roqueforti* populations grown on cheese medium or malt. However, we observed significant differences in spore production on bread medium. The Roquefort population produced the highest number of spores and significantly more than the non-Roquefort population (Fig. 5, Supplementary Table 4).

High salt concentrations have long been used in cheesemaking to prevent the growth of spoiler and pathogenic microorganisms. We found that the ability to grow on salted malt and cheese media decreased in all *P. roqueforti* populations (Supplementary Table 4). We found a significant interaction between salt and population factors, and post hoc tests indicated that the Roquefort population was more affected by salt than the other populations (Supplementary Fig. S5, Supplementary Table 4).

Volatile compound production was also investigated in the two cheese populations, as these compounds are important for cheese flavor (McSweeney, 2004). We identified 52 volatile compounds, including several involved in cheese aroma properties, such as ketones, free fatty acids, sulfur compounds, alcohols, aldehydes, pyrazines, esters, lactones and phenols (Curioni and Bosset, 2002) (Fig. 6). The two cheese populations presented significantly different volatile compound profiles, differing by three ketones, one alcohol and two pyrazines (Fig. 6). The Roquefort population produced the highest diversity of volatile compounds (Fig. 6A).

**Figure 6:**
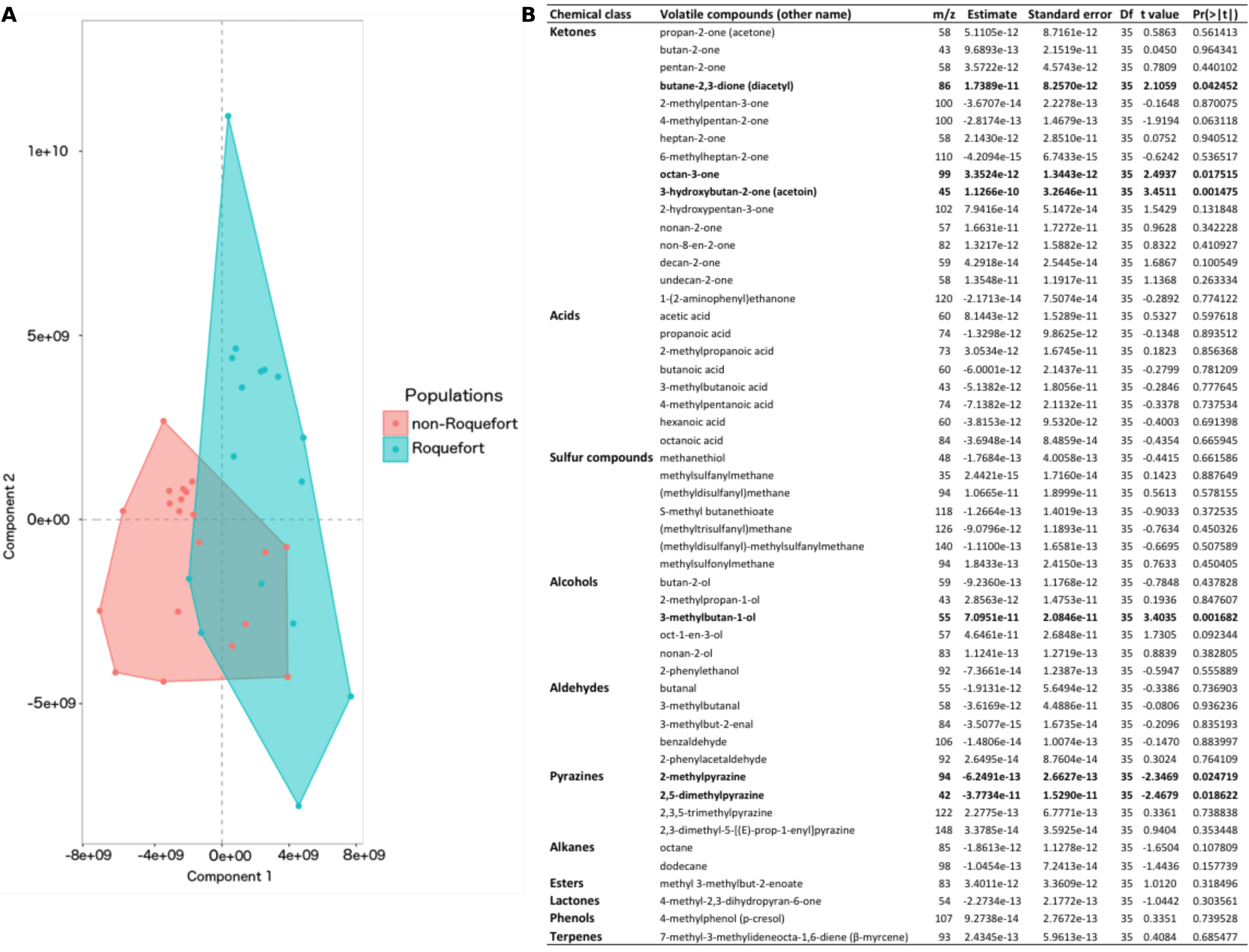
A: Differences in volatile compound profiles of the two *Penicillium roqueforti* cheese populations. Orthogonal projection of the latent structure discriminant analysis (OPLS-DA), with each dot representing the score of the averaged volatile profile of a strain from the non-Roquefort cheese population (in red) or the Roquefort population (in blue) in the two principal components. **B: Identified volatile compounds emitted by the non-Roquefort and the Roquefort populations**, chemical class, quantification ion: mass (m) to charge (z) ratio, and results of t-test statistical comparisons between the two populations: quantification estimate, standard error, degrees of freedom (Df), t values and P values (Pr(>|t|)). In bold are the volatile compounds whose quantity was found significantly different between the two populations.

### Detection of genomic regions population specific or affected by recent positive selection

We identified five regions present in the genomes of strains from the non-Roquefort cheese population and absent from the other populations. We also detected five other genomic islands present in several *P. roqueforti* strains but absent from the non-Roquefort cheese strains (Fig. 7). Nine of these ten genomic regions were not found in the genomes of the outgroup *Penicillium* species analyzed here and they displayed no genetic diversity in *P. roqueforti*. No SNPs were detected even at synonymous sites or in non-coding regions, suggesting recent acquisitions, by horizontal gene transfer. The absence of the genomic islands in some populations and outgroups prevented running gene topology analyses designed for horizontal gene transfer analyses but were even stronger evidence for the existence of horizontal gene transfer. Only FM164-C, one of the genomic islands specific to the non-Roquefort population, was present in the outgroup genomes, in which it displayed variability, indicating a loss in the other lineages rather than a gain in the non-Roquefort population and the outgroup species (Fig. 7A). The closest hits in the NCBI database for genes in the ten genomic islands were in *Penicillium* genomes. Most of the putative functions proposed for the genes within these genomic regions were related to lipolysis, carbohydrate or amino-acid catabolism and metabolite transport. Other putative functions concerned fungal development, including spore production and hyphal growth (Fig. 7). In the genomic regions specific to the non-Roquefort cheese population, we also identified putative functions potentially relevant for competition against other microorganisms, such as phospholipases, proteins carrying peptidoglycan- or chitin-binding domains and chitinases (Fig. 7) (Gooday et al., 1992). Enrichment tests were non-significant, probably due to the small number of genes in these regions.

**Figure 7:**
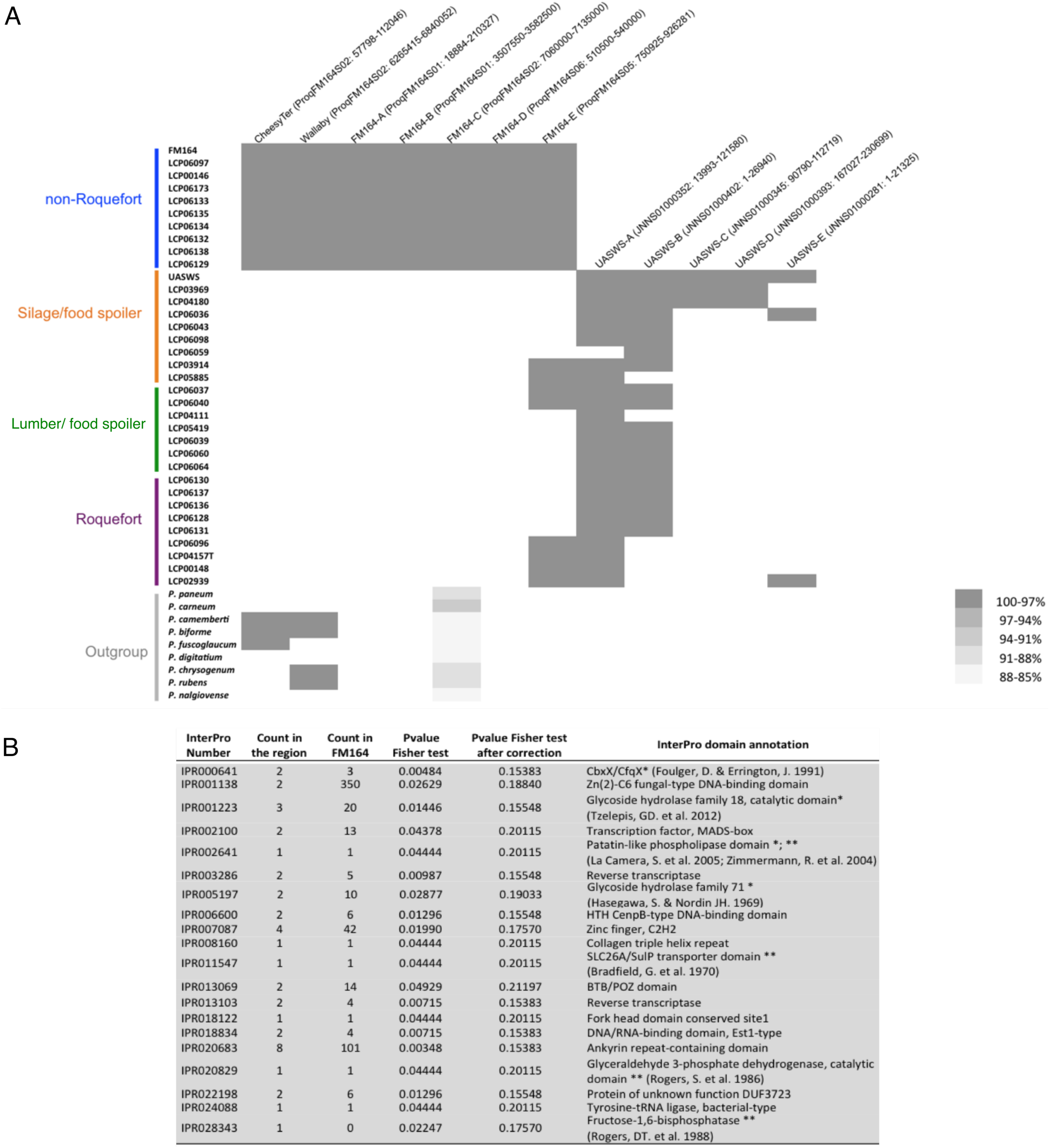
A: Presence/absence of the ten genomic islands identified in this study in the 35 *Penicillium roqueforti* and nine *Penicillium* outgroup species, in addition to the *CheesyTer* and *Wallaby* horizontally-transferred regions identified in a previous study. The ten genomic islands were detected as absent from one of the two *P. roqueforti* genomes with high-quality assemblies, while present in the second reference genome; the two reference *P. roqueforti* genomes are those of the FM164 strain (isolated from Gorgonzola cheese) and of the UASWS strain (isolated from bread Supplementary Table1 for information on outgroup reference genomes. For each genomic island, its name is indicated, together with its scaffold or contig and its start/end positions. Each strain is represented as a line, the presence of a genomic island is indicated by a colored box and its absence by a white box. The grey intensity indicates the percentage of sequence identity in these genomic islands, either within *P. roqueforti* or compared to outgroups. Strain assignment to the identified genetic clusters is indicated, with the same colors as in other figures. **B:** Fisher exact test for function enrichment of the genes identified in the presence/absence regions based on the InterPro annotation. For each annotation, the Table gives the InterPro number, the number of occurrences in the presence/absence regions and in the FM164 reference genome, the p-value before and after FDR correction and the functional annotation. Annotations are shown only for genes with significant enrichment before multiple testing correction. Annotations followed by a star refer to putative functions related to fungal growth and sporulation. Annotations followed by two stars refer to putative functions related lipolysis, carbohydrate or amino-acid catabolism and metabolite transporter.

Footprints of positive selection in *P. roqueforti* genomes were first detected using an extension of the McDonald-Kreitman test which identifies genes with more frequent amino-acid changes than expected under neutrality, neutral substitution rates being assessed by comparing the rates of synonymous and non-synonymous substitutions within and between species or populations to account for gene-specific mutation rates. We ran the test with three levels of population subdivision. First, no significant footprint of positive selection was detected for any gene by comparing the whole *P. roqueforti* species with *P. paneum*. In a second test, a set of 15 genes was identified as evolving under positive selection in the Roquefort cheese population but not in the other pooled *P. roqueforti* populations (Fig. 8A). Interestingly, eight of these 15 genes clustered at the end of the largest scaffold (Fig. 8B). In a third test, four genes were identified as evolving under positive selection in the non-Roquefort cheese population but not in the pooled non-cheese *P. roqueforti* populations (Fig. 8A). Two of these genes corresponded to a putative aromatic ring hydroxylase and a putative cyclin evolving under purifying selection in Roquefort and non-cheese *P. roqueforti* populations (Fig. 8A). Aromatic ring hydroxylases are known to be involved in the catabolism of aromatic amino acids, which are precursors of flavor compounds (Ardö, 2006; Yvon and Rijnen, 2001).

**Figure 8:**
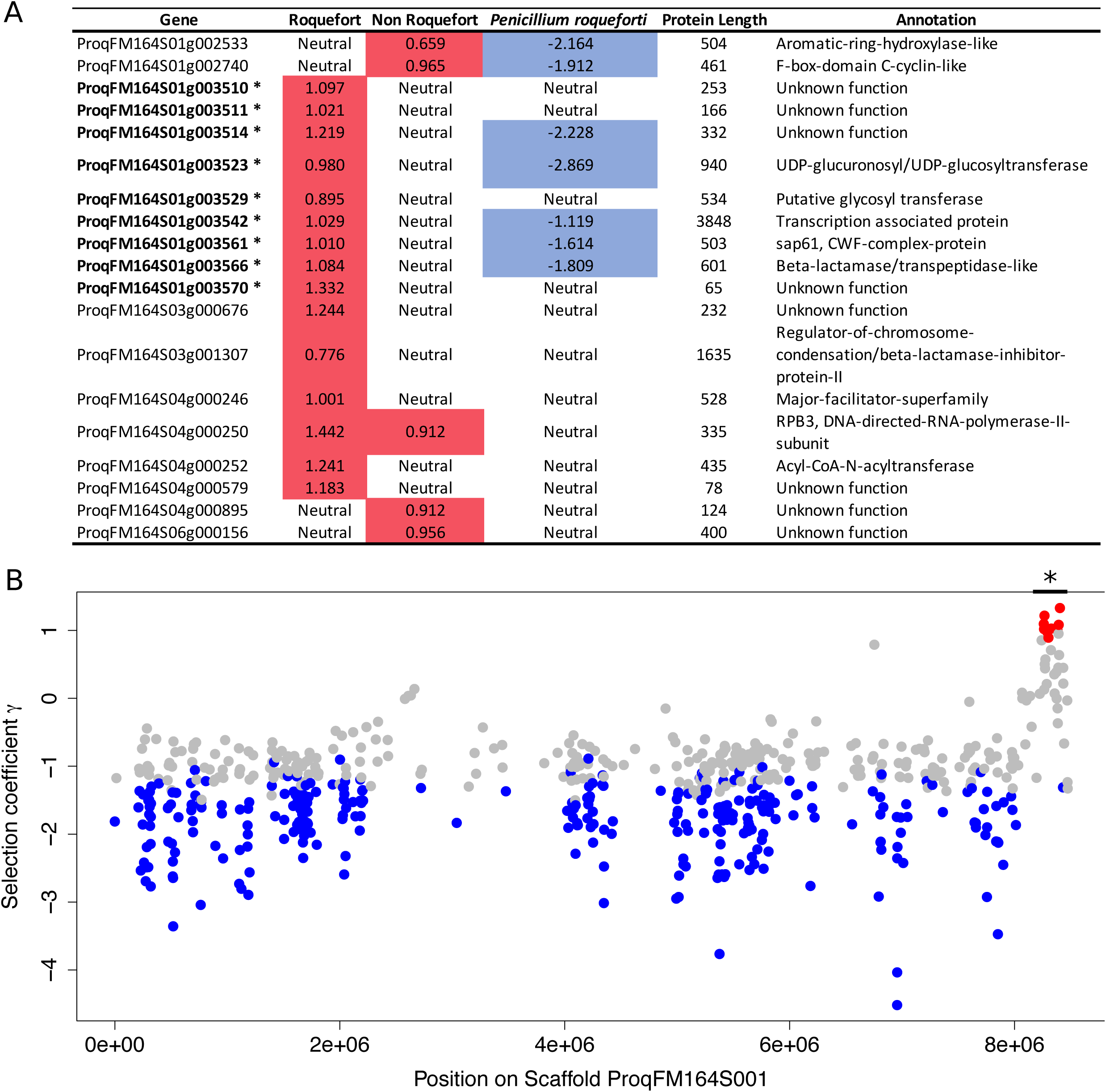
A: Genes detected as evolving under positive selection using the SnIPRE software. (i.e. genes with higher numbers of non-synonymous substitutions than expected under neutrality, controlling for gene-specific mutation rates). Values represent the estimates of the γ selection coefficient. In red, genes under positive selection (γ > 0), in blue genes under purifying selection (γ < 0), as detected based on analyses in the Roquefort cluster, the non-Roquefort cluster and in the pooled *Penicillium roqueforti* strains from the four clusters. The asterisks after the gene names highlight the eight genes clustered in the ProqFM164S01 scaffold in B. **B:** Selection effect (γ) estimated per gene along the ProqFM164S01 scaffold in the Roquefort population. The selection coefficient γ was calculated with SnIPRE. The red dots correspond to genes evolving under positive selection (γ significantly greater than 0), the blue dots to genes evolving under purifying selection (γ significantly **lower** than 0), and the gray dots to genes evolving under neutrality (γ not significantly different from 0).

Secondly, we looked for regions of low diversity and high divergence between the two cheese populations as these are footprints of recent divergent selection, i.e. positive selection in one or both of the two cheese populations but for differentiated alleles. The identified regions showed a good overlap with those detected in the Snipre analysis (Fig. 9); in particular, the same genomic island at the end of scaffold 1 stood out (Fig. 9). In the regions of high divergence and low diversity, we found a significant enrichment in transcription related genes (GO:0000981 RNA polymerase II transcription factor activity, sequence-specific DNA binding; Fisher’s exact test p-value<0.01; Supplementary Fig. 6). We found a particularly high divergence on the gene coding for RPB2 subunit of RNA polymerase II with a high number of fixed differences that were specific to the Roquefort population; fixed differences were synonymous, suggesting that important changes concern rather the regulation level than the protein itself.

**Figure 9:**
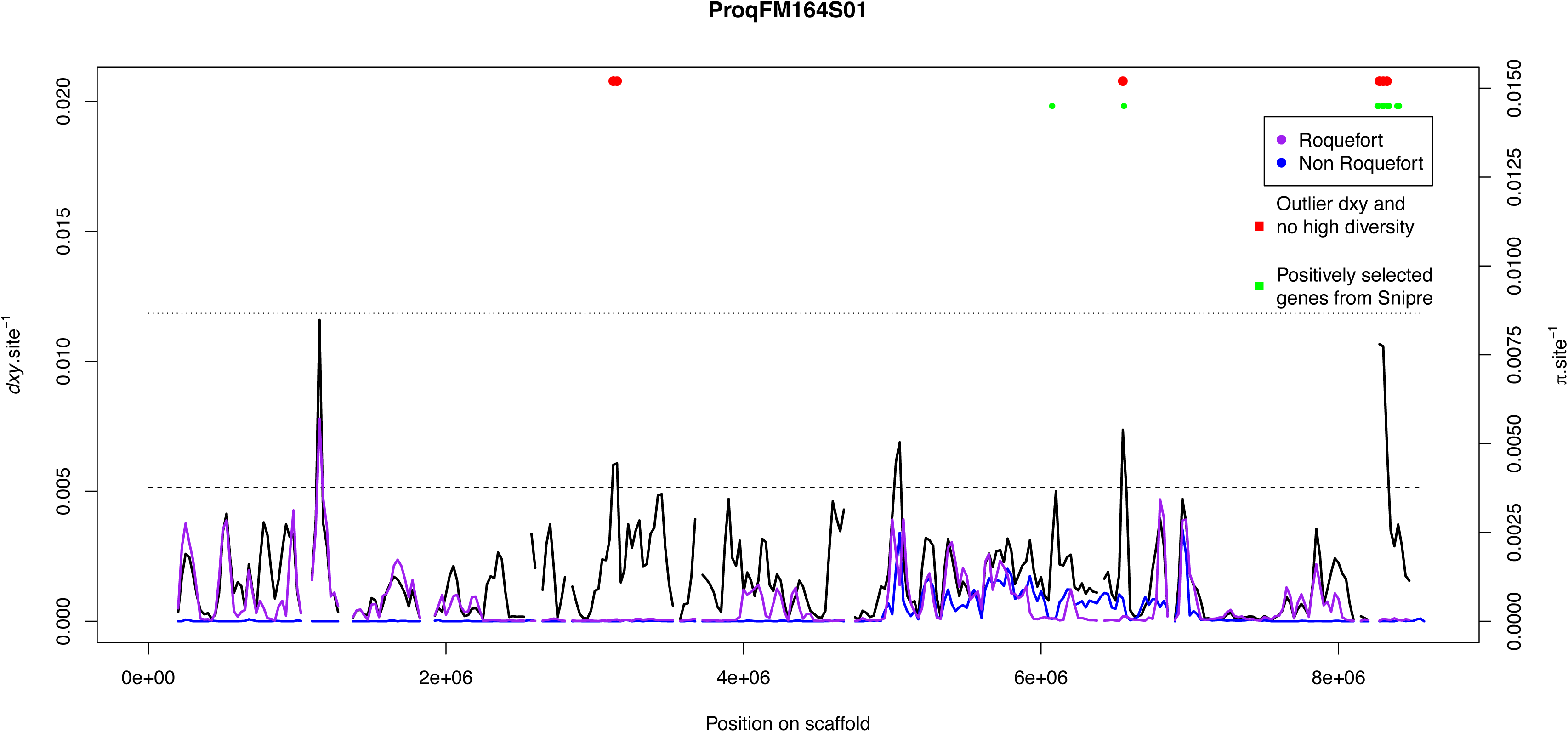
Scans of genetic differentiation (*d_xy_*) between non-Roquefort and Roquefort *Penicillium roqueforti* populations, and of genetic diversities (π) within non-Roquefort and Roquefort populations. Values were calculated in 50 kb sliding windows, overlapping over 25 kb. Red dots correspond to windows located in the 1% highest *d_xy_* (small dashed line) and 5 % lowest π values (long dashed line). Outliers detected in Snipre (Fig 8) are shown as green dots.

## Discussion

We report here the genetic subdivision of *P. roqueforti*, the fungus used worldwide for blue cheese production, with unprecedented resolution, providing insights into its domestication history. Population genomics studies on strains from various substrates and from a large collection of cheeses identified four genetically differentiated populations, two of which being cheese populations likely originating from independent and recent domestication events. One *P. roqueforti* cheese population included all the genotyped strains but one used for PDO Roquefort cheeses, produced in the French town of Roquefort-sur-Soulzon, where blue cheeses have been made since at least the 15th century, and probably long before (Aussibal, 1983; Labbe and Serres, 2009, 2004; Marre, 1906; Marres, 1935; Vabre, 2015, 2010). The strains from this Roquefort cheese population lack the horizontally-transferred *Wallaby* and *CheesyTer* genomic islands contrary to the other non-Roquefort cheese population.

We observed that the two *P. roqueforti* cheese populations differed on several traits important for cheese production, probably corresponding to historical differences. Indeed, the Roquefort population has retained moderate genetic diversity, consistent with soft selection during pre-industrial times on multiple farms near Roquefort-sur-Soulzon, where specific strains were kept for several centuries. The Roquefort cheese population grew slower in cheese (Ropars et al., 2015) and had weaker lipolytic activity. Slow maturation is particularly crucial for the storage of Roquefort cheeses for long periods in the absence of refrigeration (Marre, 1906) because they are made of ewe’s milk, a product available only between February and July. During storage, cheeses could become over degraded by too high rates of lipolysis, thus likely explaining the low lipolysis activity in Roquefort strains. By contrast, most other blue cheeses are produced from cow’s milk, which is available all year. The Roquefort population showed greater sporulation on bread, which is consistent with unconscious selection for this trait when strains were cultured on bread in Roquefort-sur-Soulzon farms before cheese inoculation during the end of the 19th and beginning of the 20th centuries.

Lipolytic activity is known to impact texture and the production of volatile compounds affecting cheese pungency (Alonso et al., 1987; De Llano et al., 1992, 1990; Martín and Coton, 2016; Thierry et al., 2017; Woo and Lindsay, 1984). The Roquefort and non-Roquefort populations showed different volatile compound profiles, suggesting also different flavor profiles. The discovery of different phenotypes in the two cheese populations, together with the availability of a protocol for inducing sexual reproduction in *P. roqueforti* (Ropars et al., 2014), pave the way for crosses to counteract degeneration after clonal multiplication and bottlenecks, for variety improvement and the generation of diversity.

Both cheese populations were found to have gone through bottlenecks. The cheese populations were the easiest to sample compared to other environments, where *P. roqueforti* is relatively rarely found. It seems therefore highly unlikely that the lower genetic diversity in the cheese populations would reflect sampling biases. In particular, the least diverse cheese population was the one including the highest numbers of countries and sampled cheese types, indicating genuine strong bottleneck. There was no particular sampling bias regarding geography either (Table S1). A previous study showed that these bottlenecks, together with clonal multiplication, decreased fertility, with different stages in sexual reproduction affected in the two populations identified here as the Roquefort and non-Roquefort lineages (Ropars et al., 2016b). The non-Roquefort cheese population, despite suffering from a more severe and more recent bottleneck, was found to be used in the production of all types of blue cheese worldwide, including Gorgonzola, Bleu d’Auvergne, Stilton, Cabrales and Fourme d’Ambert. The non-Roquefort cheese population grows more rapidly on cheese (Ropars et al., 2015), exhibits greater ability to colonize cheese cavities, higher salt tolerance and faster lipolysis than the Roquefort population. These characteristics are consistent with the non-Roquefort cheese population resulting from a very recent strong selection of traits beneficial for modern and accelerated production of blue cheese using refrigeration techniques, followed by a worldwide dissemination for the production of all types of blue cheeses. Such drastic losses of genetic diversity in domesticated organisms are typical of strong selection for industrial use by a few international firms and raise concerns about the conservation of genetic resources, the loss of which may hinder future innovation. More generally in crops, the impoverishment in genetic diversity decreases the ability of cultivated populations to adapt to environmental and biotic changes to meet future needs (Gouyon et al., 2010; Harlan, 1992; Vavilov, 1992). The PDO label, which imposes the use of local strains, has probably contributed to the conservation of genetic diversity in the Roquefort population (see “Cahier des charges de l’appellation d’origine protégée Roquefort”, i.e., the technical specifications for Roquefort PDO). We inferred two bottlenecks in the Roquefort population, more ancient than in the non-Roquefort population, likely corresponding to a pre-industrial domestication event when multiple local farms multiplied their strains, followed by a second bottleneck when fewer strains were kept by the first industrial societies. For other blue cheeses, even if their production was also ancient, the performant non-Roquefort clonal lineage could have been recently chosen to fit modern industrial production demands due to the lack of PDO rules imposing the use of local strains. However, despite a much lower genome-wide diversity in domesticated populations, proteolysis and volatile compounds diversity was found higher in cheese than in non-cheese populations. In fact, different strains with more or less rapid proteolysis and lipolysis are sold for specific blue cheese types (e.g., milder or stronger), in particular by the French LIP company (https://www.lip-sas.fr/index.php/nos-produits/penicillium-roquefortii/18-penicillium-roquefortii). Such a high phenotypic diversity within the cheese populations is consistent with diversification of usage under domestication, and in particular when different characteristics are desired according to cheese type. This has already been observed in relation to the diversification of crop varieties or breeds in domesticated animals (Parker et al., 2017; Shapiro et al., 2013).

When studying adaptation in domesticated organisms, it is often useful to contrast traits and genomic variants between domesticated and closely related wild populations to determine the nature of the adaptive changes occurring under artificial selection (Swanson-Wagner et al., 2012; Xue et al., 2016). The only known non-cheese populations of *P. roqueforti* occur essentially in human-made environments (silage, food and lumber), consistent with the specific adaptation of these populations to these environments. The two non-cheese populations were inferred to have diverged very recently, and displayed footprints of recombination and marked differentiation from the cheese populations. Domesticated populations are expected to be nested within their source populations, suggesting that we have not sampled the wild population that is the most closely related from cheese strains yet. The high level of diversity and inferred demographic history of *P. roqueforti* indicate that most food-spoiling strains belong to differentiated populations and are not feral cheese strains. In addition, not a single cheese strain was found in the food spoiling and silage populations. This was shown by both genome sequences and by the genotyping of a larger number of strains using a few selected markers, in the present study and based on microsatellite markers in a previous work (Ropars et al., 2017). Consequently, *P. roqueforti* spores from blue cheeses may, rarely, spoil food and food-spoiling and silage strains are not used for cheesemaking nor recombine with cheese strains. Such a lack of incoming gene flow into cheese populations allowed trait differentiation in cheese strains as expected under domestication.

It came as a surprise that the two non-cheese populations split more recently from each other than from the cheese lineages. In particular, the non-Roquefort population diverged the earliest from the unidentified ancestral population, and this has likely occurred in another environment than cheese. Much more recently, selection in industrial times has likely only kept the most performant clonal lineage of this population for cheesemaking, losing most of the initial diversity as indicated by the very strong and recent bottleneck inferred in this lineage. Possible scenarios to explain the existence of two separated clusters thriving in food and silage differentiated from cheese strains include the very recent adaptive differentiation of a population from silage on human food or vice versa. The finding that silage strains are only found in one cluster (the orange one in Fig.1 to 5) suggests an adaptation to this ecological niche, although experiments will be required to test this hypothesis. Food spoiling strains are in contrast found in three clusters and may thus not constitute a specific population adapted to this environment and may instead represent migrants from several populations belonging to other ecological niches. Green and orange clusters may alternatively represent populations thriving in yet unidentified environments, dispersing to silage and food. Another hypothesis would be a single domestication event for cheesemaking before the divergence of the four lineages, followed by an escape and subsequent differentiation of the orange and green lineages in other human related habitats. This hypothesis however would not predict such high genetic diversity in the green and orange populations, and in particular the similar nucleotidic diversity levels in the two non-cheese populations as in the *P. carneum* and *P. paneum* outgroups. Given the very low genetic diversity in the cheese populations, coalescence events occurred recently in the past, preventing tests of the occurrence of bottlenecks in the common ancestor of the four *P. roqueforti* populations.

The history of blue cheese production may provide circumstantial clues as to the origin of *P. roqueforti* cheese populations. Indeed, the first blue cheeses likely resulted from the sporadic accidental contamination of cheese with spores from the environment, such as moldy food. However, this would not be consistent with the demographic history inferred here for cheese and food-spoiling strains, as the cheese strains were not found to be nested within the food-spoiling strains, some of which originated from moldy bread. Furthermore, old French texts suggest that the blue mold colonized the cheese from within (Labbe and Serres, 2009, 2004; Vabre, 2015), which would indicate that the milk or curd was contaminated. French cheese producers began to inoculate cheeses with *P. roqueforti* spores from moldy rye bread at the end of the 19^th^ century (Labbe and Serres, 2009, 2004; Vabre, 2015). Breads were specifically made with a 2:1 mixture of wheat and rye flour and were baked rapidly at high temperature (500°C), to yield a protective crust, around a moist, undercooked interior (Aussibal, 1983; Marre, 1906); the mold developed from the inside of the bread after one to five months in the Roquefort caves (Labbe and Serres, 2009, 2004; Vabre, 2015). Surveys of the microorganisms present in their caves (Chaptal, 1789; Marcorelle and Chaptal, 1833; Marre, 1906) and our unsuccessful attempts to obtain samples from a maturing cellar suggest that *P. roqueforti* spores did not originate from the caves, which were nevertheless crucial due to the ideal conditions provided for *P. roqueforti* development (Marre, 1906). Bread may have been colonized from the environment or from rye flour if the source *P. roqueforti* population was a rye endophyte or pathogen. This last hypothesis would be consistent with the lifestyle of many *Penicillium* species, which live in close association with plants, often acting as plant pathogens or necrotrophs (Ropars et al., 2016a), and with the occurrence of a *P. roqueforti* population in lumber and silage. Actually, a recent study reports the finding of *P. roqueforti* as an endophyte and could be inoculated on wheat (Ikram et al., 2018), although species identification should be checked with more powerful markers. If this hypothesis is correct, then cheeses may historically have become contaminated with *P. roqueforti* from fodder during milking.

Comparison between non-cheese and cheese populations allowed us to identify specific traits and genes that have been under selection in cheese as opposed to other environments. Furthermore, the two independently domesticated *P. roqueforti* cheese populations, exhibiting different traits, represent a good model for studying the genomic processes involved in adaptation. We could not run analyses of selective sweep detection based on local decrease in genetic diversity in the genomes; indeed, because of the clonality of cheese populations, the whole genome will have hitchhiked with any selected locus. This effect has likely contributed to the strong bottlenecks. We were nevertheless able to identify candidate genes and evolutionary mechanisms potentially involved in adaptation to cheese in *P. roqueforti*. The horizontally-transferred *CheesyTer* genomic island probably contributes to the faster growth of the strains identified here as constituting the non-Roquefort population (Ropars et al., 2015). Indeed, *CheesyTer* includes genes with putative functions involved in carbohydrate utilization (e.g. β-galactosidase and lactose permease genes) that are specifically expressed at the beginning of cheese maturation, when lactose and galactose are available. This horizontal gene transfer may thus have been involved in adaptation to recently developed industrial cheese production processes in the non-Roquefort cheese population, conferring faster growth. We also identified additional genomic islands specific to the non-Roquefort cheese population, probably acquired recently and including genes putatively involved in fungal growth and spore production. In the genomic islands specific to the cheese populations, several genes appeared to be involved in lipolysis, carbohydrate or amino-acid catabolism and metabolite transport, all of which are important biochemical processes in the development of cheese flavor. In the Roquefort cheese population, a genomic region harboring genes with footprints for positive selection included several genes encoding proteins potentially involved in aromatic amino-acid catabolism corresponding to precursors of volatile compounds. Further studies are required to determine the role of these genes in cheese flavor development.

In conclusion, we show that *P. roqueforti* cheese populations represent genuine domestication. Of course, the domestication process in cheese fungi has been more recent and different from the ones in emblematic crops or animals. Nevertheless, we did observe strong genetic differentiation from non-cheese populations, strong bottlenecks and trait differentiation with likely benefits for cheese production. This suggests genuine domestication, as has been reported previously in other fungi (Almeida et al., 2014; Baker et al., 2015; Gallone et al., 2016; Gibbons et al., 2012; Gonçalves et al., 2016; Libkind et al., 2011; Sicard and Legras, 2011), and defined as “the genetic modification of a species by breeding it in isolation from its ancestral population in an effort to enhance its utility to humans” (Gibbons and Rinker, 2015). Furthermore, a previous study has shown that the non-Roquefort cheese strains have acquired genes conferring better growth in cheese (Ropars et al., 2015). Our study revealed genetic divergence of cheese population from non-cheese populations, as well as the evolution of specific traits, with beneficial characteristics for cheese production. These findings therefore indicate the occurrence of domestication, a special case of adaptive divergence. We found that gene flow was prevented by clonality of cheese lineages and lack of migration between cheese and non-cheese populations, and that adaptation occurred on several traits beneficial for cheese production (lipolysis, proteolysis, spore production, volatile compound production, growth in salted cheese, cheese cavity colonization ability). Genomic footprints of adaptation were found in terms of rapid amino-acid changes and horizontal gene transfers. The two independent domestication events identified here interestingly represent adaptations to different production modes. Our findings concerning the history of *P. roqueforti* domestication thus shed light on the processes of adaptation to rapid environmental change, but they also have industrial implications and raise questions about the conservation of genetic resources in the agri-food context.

## Methods

### Isolation attempts of Penicillium roqueforti in ripening cellar and dairy environments

In order to investigate whether a wild *P. roqueforti* population occurred in ripening cellars or dairy environments that could be at the origin of the observed cheese populations, we sampled spores from the air in an artisanal cheese dairy company (GAEC Le Lévejac, Saint Georges de Lévejac, France, ca 60 km from Roquefort-sur-Soulzon, producing no blue cheese to avoid feral strains, i.e. dispersal from inoculated cheeses), sampling was performed in the sheepfold, milking parlour, cheese dairy and ripening cellar. We also sampled spores from the air in an abandoned ripening cellar in the town of Meyrueis (ca 70 km from Roquefort-sous-Soulzon) where Roquefort cheeses used to be produced and stored in the early 19^th^ century. In total, 55 Petri dishes containing malt (2% cristomalt, Difal) and 3% ampicillin were left open for six days as traps for airborne spores (35 Petri dishes in the abandoned ripening cellar and 20 Petri dishes in the artisanal cheese dairy company). Numerous fungal colonies were obtained on the Petri dishes. One monospore was isolated from each of the 22 *Penicillium*-like colonies. DNA was extracted using the Nucleospin Soil Kit (Macherey-Nagel, Düren, Germany) and a fragment of the β-tubulin gene was amplified using the primer set Bt2a/Bt2b (Glass and Donaldson, 1995), and then sequenced. Sequences were blasted against the NCBI database to assign monospores to species. Based on β-tubulin sequences, ten strains were assigned to *P. solitum*, six to *P. brevicompactum*, two to *P. bialowienzense*, one to *P. echinulatum* and two to the *Cladosporium* genus. No *P. roqueforti* strain could thus be isolated from this sampling procedure.

### Genome sequencing and analysis

The genomic DNAs of cheesemaking strains obtained from public collections belonging to *P. roqueforti*, seven strains of *P. paneum,* one strain of *P. carneum* and one strain of *P. psychrosexualis* (Supplementary Table 1) were extracted from fresh haploid mycelium after monospore isolation and growth for five days on malt agar using the Nucleospin Soil Kit (Macherey-Nagel, Düren, Germany). Sequencing was performed using the Illumina HiSeq 2500 paired-end technology (Illumina Inc.) with an average insert size of 400 bp at the GenoToul INRA platform and resulted in a 50x-100x coverage. In addition, the genomes of four strains (LCP05885, LCP06096, LCP06097 and LCP06098) were used that had previously been sequenced using the ABI SOLID technology (Cheeseman et al., 2014). GenBank accession numbers are HG792015-HG792062.

Identification of presence/absence polymorphism of blocks larger than 10 kbp in genomes was performed based on coverage using mapping against the FM164 *P. roqueforti* reference genome. In order to identify genomic regions that would be lacking in the FM164 genome but present in other strains, we used a second assembled genome, that of the UASWS *P. roqueforti* strain collected from bread, sequenced using Illumina HiSeq shotgun and displaying 428 contigs (Genbank accession numbers: JNNS01000420-JNNS01000428). Blocks larger than 10 kbp present in the UASWS genome and absent in the FM164 genome were identified using the *nucmer* program v3.1 (Kurtz et al., 2004). Gene models for the UASWS genome were predicted with EuGene following the same pipeline as for the FM164 genome (Cheeseman et al., 2014; Foissac et al., 2008). The presence/absence of these regions in the *P. roqueforti* genomes was then determined using the coverage obtained by mapping reads against the UASWS genome with the start/end positions identified by *nucmer*. The absence of regions was inferred when less than five reads were mapped. In order to determine their presence/absence in other *Penicillium* species, the sequences of these regions were blasted against nine *Penicillium* reference genomes (Supplementary Table 1). PCR primer pairs were designed using Primer3Plus (http://www.bioinformatics.nl/cgi-bin/primer3plus/primer3plus.cgi/) in the flanking sequences of these genomic regions in order to check their presence/absence in a broader collection of *P. roqueforti* strains based on PCR tests (Supplementary Table 2). For each genomic island, two primer pairs were designed when possible (i.e. when sufficiently far from the ends of the scaffolds and not in repeated regions): one yielding a PCR product when the region was present and another one giving a band when the region was absent, in order to avoid relying only on lack of amplification for inferring the absence of a genomic region. PCRs were performed in a volume of 25 µL, containing 12,5 µL template DNA (ten folds diluted), 0.625 U Taq DNA Polymerase (MP Biomedicals), 2.5 µL 10x PCR buffer, 1 µL of 2.5 mM dNTPs, 1 µL of each of 10 µM primer. Amplification was performed using the following program: 5 min at 94°C and 30 cycles of 30 s at 94°C, 30 s at 60°C and 1 min at 72°C, followed by a final extension of 5 min at 72°C. PCR products were visualized using stained agarose gel electrophoresis. Data were deposited at the European Nucleotide Archive (http://www.ebi.ac.uk/ena/) under the accession number: PRJEB20132 for whole genome sequencing and PRJEB20413 for Sanger sequencing.

For each strain, reads were mapped using stampy v1.0.21 (Lunter and Goodson, 2011) against the high-quality reference genome of the FM164 *P. roqueforti* strain (Cheeseman et al., 2014). In order to minimize the number of mismatches, reads were locally realigned using the genome analysis toolkit (GATK) IndelRealigner v3.2-2 (McKenna et al., 2010). SNP detection was performed using the GATK Unified Genotyper (McKenna et al., 2010), based on the reference genome in which repeated sequences were detected using RepeatMasker (Smit et al., 2013) and masked, so that SNPs were not called in these regions. In total 483,831 bp were masked, corresponding to 1.67% of the FM164 genome sequence. The 1% and 99% quantiles of the distribution of coverage depth were assessed across each sequenced genome and SNPs called at positions where depth values fell in these extreme quantiles were removed from the dataset. Only SNPs with less than 10% of missing data were kept. After filtering, a total of 115,544 SNPs were kept.

Population structure was assessed using a discriminant analysis of principal components (DAPC) with the Adegenet R package (Jombart, 2008). The genetic structure was also inferred along the genome by clustering the strains according to similarities of their genotypes, in windows of 50 SNPs, using the Mclust function of the mclust R package (Fraley et al., 2012; Fraley and Raftery, 2002) with Gower’s distance and a Gaussian mixture clustering with K=7 (as the above analyses indicated the existence of four *P. roqueforti* populations and there were three outgroup species).

We performed a neighbor-net analysis using the network approach to visualize possible recombination events within and between populations with the phangorn R package (Schliep, 2010). The substitution model used for building the distance matrix was JC69 (Jukes and Cantor, 1969)

The genetic diversity were estimated using the θπ and, θw with the compute programs associated to libsequence v1.8.9 (Thornton, 2003) on 1145 sliding windows of 50 kb with 25 kb of overlap distributed along the longest eleven scaffolds of the FM164 assembly (> 200 kb). Linkage disequilibrium per genetic cluster (i.e. non-Roquefort, Roquefort, Lumber/food spoiler and silage/food spoiler) was estimated using the r2 statistics, with VCFtools v 0.1.15 (Danecek et al., 2011) and the following parameters: --geno-r2 --ld-window-bp 15000. Plots were generated using R.

To identify genes evolving under positive selection in *P. roqueforti* genomes, first, we used the method implemented in SnIPRE (Eilertson et al., 2012), a Bayesian generalization of the log-linear model underlying the McDonald-Kreitman test. This method detects genes in which amino-acid changes are more frequent than expected under neutrality, by contrasting synonymous and non-synonymous SNPs, polymorphic or fixed in two groups, to account for gene-specific mutation rates. Secondly, we performed a scan of the divergence statistics dxy between the two cheese populations, calculated using a custom R script in 50kbp windows overlapping over 25 kbp along the genome. We considered genes belonging to the 1% most divergent regions and the 5% least genetically diverse (π values) as under positive selection in one of the populations. We did not consider the other pairwise comparisons, i.e. using orange and green populations (Figs. 1 to 5), because most SNPs in those populations were shared by several strains, as shown by high diversity, positive Dt and low FST values (Table 1). Consequently, islands of high divergence and low diversity were restricted to cheese populations that were already found using pairwise comparison between cheese populations. We performed GO annotation enrichment tests using separate Fisher’s exact tests on the three ontologies (BP: biological process; CC: cellular component; MF: metabolic function).

### Strain genotyping

We identified two genomic regions with multiple diagnostic SNPs allowing discriminating the two cheese clusters. Two PCR primer pairs were designed (Supplementary Table 2) to sequence these regions in order to assign the 65 strains (Supplementary Table 1) that can be purchased at the Laboratoire Interprofessionnel de Production d’Aurillac (LIP) (the main French supplier of *P. roqueforti* spores for artisanal and industrial cheese-makers; https://www.lip-sas.fr/) to the identified clusters. PCR products were then purified and sequenced at Eurofins (France). Because one of the cheese clusters included strains carrying the *Wallaby* and *CheesyTer* genomic islands while the second cluster strains lacked these genomic regions (Ropars et al., 2015), we used previously developed primer pairs to check for the presence/absence of *CheesyTer* and *Wallaby* (Ropars et al., 2015).

Sequences were first aligned together with those extracted from sequenced genomes, allowing assignation of LIP strains to one of the two cheese populations using MAFFT software (Katoh and Standley, 2013) and then the alignments were visually checked. Then a tree reconstruction was made using RAxML following GTRCAT substitution model, using 2 partitions corresponding to the two fragments and 1000 bootstraps tree were generated (Stamatakis, 2006).

### Strain phenotyping

For all experiments, strains were picked up at random in each group. Experimental cheeses were produced in an artisanal dairy company (GAEC Le Lévejac, Saint Georges de Lévejac, France). The same ewe curd was used for all produced cheeses. Seven *P. roqueforti* strains were used for inoculation (two from each of the Roquefort, non-Roquefort and silage/food spoiler clusters, and one from the lumber/food spoiler cluster; their identity is given in Supplementary Table 1) using 17.8 mg of lyophilized spores. Three cheeses were produced for each strain in cheese strainers (in oval pots with opposite diameters of 8 and 9 cm, respectively), as well as a control cheese without inoculation. After 48 h of draining, cheeses were salted (by surface scrubbing with coarse salt), pierced and placed in a maturing cellar for four weeks at 11°C. Cheeses were then sliced into six equal pieces and a picture of each slice was taken using a Nikon D7000 (zoom lens: Nikon 18-105mm f:3.5-5.6G). Pictures were analyzed using the geospatial image processing software ENVI (Harris Geospatial Solution) (Fig. 6). This software enables pixel classification according to their level of blue, red, green, and grey into two to four classes depending on the analyzed image. This classification allowed assigning pixels to two classes corresponding to the inner white part and the cavities of the cheese, respectively (Fig. 6). For each picture, the percentage of pixels corresponding to the cavities was then quantified. Because the software could not reliably assign pixels to the presence versus absence of the fungus in cavities, we visually determined the cavity areas that were colonized by *P. roqueforti* using images. This allowed calculating a cheese cavity colonization rate. Because *Penicillium* spores have a high dispersal ability which could cause contaminations, we confirmed strain identity present in cheeses by performing Sanger sequencing of four diagnostic markers designed based on SNPs and specific to each strain (Supplementary Table 2). For each cheese, three random monospore isolates were genotyped, and no contamination was detected (i.e. all the sequences obtained corresponded to the inoculated strains).

To compare the growth rates of the different *P. roqueforti* clusters on bread (i.e. the traditional multiplication medium), 24 strains were used (eight from each of the Roquefort and non-Roquefort cheese clusters, five from the silage/food spoiler cluster, and three from the lumber/food spoiler cluster; the identities of the strains are shown in Supplementary Table 1). Each strain was inoculated in a central point in three Petri dishes by depositing 10 µL of a standardized spore suspension (0.7×10^9^ spores/mL). Petri dishes contained agar (2%) and crushed organic cereal bread including rye (200 g/L). After three days at 25°C in the dark, two perpendicular diameters were measured for each colony to assess colony size.

The lipolytic and proteolytic activities of *P. roqueforti* strains were measured as follows: standardized spore suspensions (2500 spores/inoculation) for each strain (n=47: 15 from the Roquefort cluster, 15 from the non-Roquefort cheese cluster, 10 from the silage/food spoiler cluster and seven from the lumber/food spoiler cluster, identity in Supplementary Table 1) were inoculated on the top of a test tube containing agar and tributyrin for lypolytic activity measure (10 mL/L, ACROS Organics, Belgium) or semi-skimmed milk for the proteolytic activity measure (40 g/L, from large retailers). The lipolytic and proteolytic activities were estimated by the degradation degree of the compounds, which changes the media from opaque to translucent. For each media, three independent experiments have been conducted. For each strain, duplicates were performed in each experiment and the limit of translucency / opaqueness in the medium was recorded. Measures were highly repeatable between the two replicates (Pearson’s product-moment correlation coefficient of 0.93 in pairwise comparison between replicates, P<0.0001). We measured the distance between the initial mark and the hydrolysis, translucent front, after 7, 14, 21 and 28 days of growth at 20°C in the dark.

A total of 47 strains were used to compare spore production between the four *P. roqueforti* clusters (Supplementary Table 1), 15 belonging to the non-Roquefort cluster, 15 to the Roquefort cluster, 10 to the silage/food spoiler cluster and seven to the lumber/food spoiler cluster. After seven days of growth on malt agar in Petri dishes of 60 mm diameter at room temperature, we scraped all the fungal material by adding 5 mL of tween water 0.005%. We counted the number of spores per mL in the solution with a Malassez hemocytometer (mean of four squares per strain) for calibrating spore solution. We spread 50 μL of the calibrated spore solution (i.e. 7.10^6^ spores.mL^-1^) for each strain on Petri dishes of 60 mm diameter containing three different media, malt, cheese and bread agar (organic “La Vie Claire” bread mixed with agar), in duplicates (two plates per medium and per strain). After eight days of growth at room temperature, we took off a circular plug of medium with spores and mycelium at the top, using Falcon 15 mL canonical centrifuge tubes (diameter of 15 mm). We inserted the plugs into 5 mL Eppendorf tubes containing 2 mL of tween water 0.005% and vortexed for 15 seconds to detach spores from the medium. Using a plate spectrophotometer, we measured the optical density (OD) at 600 nm for each culture in the supernatant after a four-fold dilution (Supplementary Table 3).

To compare salt tolerance between *P. roqueforti* clusters, 26 strains were used (eight from the Roquefort cluster, ten from the non-Roquefort cluster, three from the silage/food spoiler cluster, and five from the lumber/food spoiler cluster; strain identities are shown in Supplementary Table 1). For each strain and each medium, three Petri dishes were inoculated by depositing 10 µL of standardized spore suspension (0.7×10^9^ spores/mL) on Petri dishes containing either only malt (20 g/L), malt and salt (NaCl 8%, which corresponds to the salt concentration used before fridge use to avoid contaminants in blue cheeses), only goat cheese, or goat cheese and salt (NaCl 8%). The goat cheese medium was prepared as described in a previous study (Ropars et al., 2015). Strains were grown at 25°C and colony size measured daily for 24 days.

Volatile production assays were performed on 16 Roquefort strains and 19 non-Roquefort cheese strains grown on model cheeses as previously described (Gillot et al., 2017). Briefly, model cheeses were prepared in Petri dishes and incubated for 14 days at 25 °C before removing three 10 mm-diameter plugs (equivalent to approximately 1 g). The plugs were then placed into 22 mL Perkin Elmer vials that were tightly closed with polytetrafluorethylene (PTFE)/silicone septa and stored at −80°C prior to analyses (Gillot et al., 2017). Analyses and data processing were carried out by headspace trap-gas chromatography-mass spectrometry (HS-trap-GC-MS) using a Perkin Elmer turbomatrix HS-40 trap sampler, a Clarus 680 gas chromatograph coupled to a Clarus 600T quadrupole MS (Perkin Elmer, Courtaboeuf, France), and the open source XCMS package of the R software (http://www.r-project.org/), respectively, as previously described (Pogačić et al., 2015).

All phenotypic measures are reported in Supplementary Table 3. Statistical analyses for testing differences in phenotypes between populations and/or media (Supplementary Table 4) were performed with R software (http://ww.r-project.org).

Differences in volatile profiles among the two *P. roqueforti* cheese populations were analyzed using a supervised multivariate analysis method, orthogonal partial least squares discriminant analysis (OPLS-DA). OPLS is an extension of principal components analysis (PCA), that is more powerful when the number of explained variables (Y) is much higher than the number of explanatory variables (X). PCA is an unsupervised method maximizing the variance explained in Y, while partial least squares (PLS) maximizes the covariance between X and Y(s). OPLS is a supervised method that aims at discriminating samples. It is a variant of PLS which uses orthogonal (uncorrelated) signal correction to maximize the explained covariance between X and Y on the first latent variable, and components >1 capture variance in X which is orthogonal (uncorrelated) to Y. The optimal number of latent variables was evaluated by cross-validation (Pierre et al., 2011). Finally, to identify the volatile compounds that were produced in significantly different quantities between the two populations, a t-test was performed using the R software (http://www.r-project.org/).

### Demographic modeling using approximate Bayesian computation (ABC)

The likelihoods of 11 demographic scenarios for the *P. roqueforti* populations were compared using approximate Bayesian computation (ABC) (Beaumont, 2010; Lopes and Beaumont, 2010). The scenarios differed in the order of demographic events, and included 21 parameters to be estimated (Supplementary Fig. 4). A total of 262 fragments, ranging from 5 kb to 15 kb, were generated from observed SNPs by compiling in a fragment all adjacent SNPs in complete linkage disequilibrium. The population mutation rate θ (the product of the mutation rate and the effective population size) used for coalescent simulations was obtained from data using θ_w,_ the Watterson’s estimator. Simulated data were generated using the same fragment number and sizes as the SNP dataset generated from the genomes. Priors were sampled in a log-uniform distribution (Supplementary Fig. 4C). For each scenario, one million coalescent simulations were run and the following summary statistics were calculated on observed and simulated data using msABC (Pavlidis et al., 2010) : the number of segregating sites, the estimators π (Nei, 1987) and θ_w_ (Watterson, 1975) of nucleotide diversity, Tajima’s D (Tajima, 1989), the intragenic linkage disequilibrium coefficient ZnS (Kelly, 1997), F_ST_ (Hudson et al., 1992), the percentage of shared polymorphisms between populations, the percentage of private SNPs for each population, the percentage of fixed SNPs in each population, Fay and Wu’s H (Fay and Wu, 2000), the number of haplotypes (Depaulis and Veuille, 1998) and the haplotype diversity (Depaulis and Veuille, 1998). For each summary statistic, both average and variance values across simulated fragments were calculated. The choice of summary statistics to estimate posterior parameters is a crucial step in ABC (Csilléry et al., 2010). Summary statistics were selected using the AS.select() function with the neuralnet method in the “abctools” R package (Nunes and Prangle, 2015). In total, 101 summary statistics were kept for subsequent analyses. Cross validation was run with the neuralnet method using 100 samples and a tolerance of 0.01 (Supplementary Fig. 4D). Model selection was performed using four tolerance rates ranging from 0.005 to 0.1 and rejection, logistic regression and neural network methods. Because there was still an uncertainty on the choice between scenarios 4 and 5 after model selection (i.e. whether it was the non-Roquefort or Roquefort population that diverged first from the ancestral population) (Supplementary Fig. 4 F), an extra one million simulations were run for each of those two scenarios and model selection was performed again. All tolerance rates and methods favored scenario 4 over scenario 5 with absolute confidence of 1.000.

The posterior probability distributions of the parameters, the goodness of fit for each model and model selection (Supplementary Fig. 4E) were calculated using a rejection-regression procedure (Beaumont, 2010). Acceptance values of 0.005 were used for all analyses. Regression analyses was performed using the “abc” R package (Csilléry et al., 2012) (http://cran.rproject.org/web/packages/abc/index.html).

### Estimate of time since domestication

The multiple sequentially Markovian coalescent (MSMC) software was used to estimate the domestication times of cheese populations (Schiffels and Durbin, 2014). The estimate of the last time gene flow occurred within each cheese population was taken as a proxy of time since domestication as it also corresponds in such methods to bottleneck date estimates and is more precisely estimated. Recombination rate was set at zero because sexual reproduction has likely not occurred since domestication in cheese populations (see results). Segments were set to 21*1+1*2+1*3 for the Roquefort population which contains three haplotypes (Fig. 2) and to 10*1+15*2 for the non-Roquefort population, which contains two closely related haplotypes ( Fig. 2). In both cases, MSMC was run for 15 iterations and otherwise default parameters. The mutation rate was set to 10^-8^.

## Acknowledgments

This work was supported by the ERC starting grant GenomeFun 309403 awarded to TG, the ANR FROMA-GEN grant (ANR-12-PDOC-0030) to AB, and an “*Attractivité*” grant from Paris-Sud University to AB. We thank Kamel Soudani for help with image analysis and Aurélien Tellier for advice concerning ABC analyses. We are grateful to Coralie Benel and Francis Roujon of GAEC Le Lévejac for assistance with cheesemaking and Paul Villain for experimental help. Sequencing was performed at GenoToul INRA platform. We thank INRA and MNHN for granting access to four genomes sequenced with the help of Joëlle Dupont, Sandrine Lacoste, Yves Brygoo and Jeanne Ropars in the framework of the ANR ‘Food Microbiomes’ project (ANR-08-ALIA-007-02) coordinated by Pierre Renault.

## Author contributions

TG and AB acquired the funding, designed and supervised the study. SL and AS produced the genomes. ED, AB and RdlV analyzed the genomes. ED, SL, JR, AS, MC, AT, EC, MLP and DR performed the experiments. ED, AB and TG analyzed the data from the experiments. ED, AB and AF performed ABC analyses. ED and TG wrote the manuscript with contributions from the other authors.

## Notes

#### Summary of Updates

Figures updated Text updated Some new analyses

