## Supplementary Legends and Figures for "Independent domestication events in the blue-cheese fungus *Penicillium roqueforti*"

### Supplementary Inventory

#### Supplementary Figure 1

**Decay of linkage disequilibrium (LD) in each *Penicillium roqueforti* population.** The $r^2$ parameters was calculated pairwise between all SNPs that are on the six largest scaffolds with a maximum distance of 80,000 bp. The average $r^2$ for each inter-SNPs distance was calculated and used for plotting.

#### Supplementary Figure 2

**Population structure analysis** using the fastSTRUCTURE program with number of clusters *K* ranging from 2 to 6. Each individual is represented by a vertical bar divided into *K* colored segments representing the likelihood of membership to the corresponding clusters. The marginal likelihood is reported for each K tested.

#### Supplementary Figure 3

**Consensus tree at 50% from 100 bootstrap replicates of the *Penicillium roqueforti* strains from different sources based on concatenated sequences of two genes** The strains isolated or used for Roquefort and other blue cheese clusters are labelled in blue and purple, respectively. Strains which whole genome has been sequenced are label with a (*). Support values are derived from 1,000 bootstrap replicates. The scale bar indicates the number of substitutions. Strains either come from the LCP (laboratoire de cryptogamie) collection belonging to the National Museum of Natural History in Paris, from the LIP collection (Laboratoire Interprofessionnel de Production d’Aurillac), the main French supplier of *P. roqueforti* spores for cheesemakers or from the UBOCC collection of the Laboratoire de Biodiversité et d’Ecologie Microbienne at Université de Bretagne Occidentale.

#### Supplementary Figure 4

**A Approximate Bayesian computation (ABC) scenarios.** The colors denote the four populations as in other figures. Schematic representation of the eleven demographic scenarios for the history of *Penicillium roqueforti* populations, compared using ABC; **B Demographic parameters shown on scenario S4 as an example**: t=time in unit of $2N_e$; m=migration rate; g=growth rate; n=ancestral population size reduction coefficient. **C Definition and prior distribution of parameters used in the approximate Bayesian computation (ABC) tests of scenarios.** Example of scenario S4 (Constraints on parameters : t8<t11 t10<t8; t9<t10; t3<t8; t12<t3; t2<t12; t1<t2; t5<t12; t4<t5; t7<t11; t6<t7). **D Confusion matrix for the eleven scenarios tested. E**ach number on x-axis and each color represent a scenario described in A. For each scenario, 1000 samples of cross validation were simulated to estimate the power to estimate the good scenario from our data. In each bar, each color represents the proportion of simulations where the scenario of the corresponding color was recovered from the neural network method using a tolerance rate of 0.01 and with the set of chosen summary statistics. **E Parameter estimates for the scenario S4.** Parameters were inferred from the posteriors obtained with the S4 model. For each parameter, the 95% credible interval of the mean *a posteriori* estimate is given (in brackets) using rejection algorithm. **F Approximate Bayesian computation (ABC) comparison between the eleven evolutionary scenarios and three tolerance rates. P**osterior model probabilities using the two regression methods « Mnlogistic » (where the posterior model probabilities are estimated using a multinomial logistic regression) and « Neuralnet » (where neural networks are used to predict the probabilities of models based on the observed statistics). Rejection represents the proportion of accepted simulations among the 1 million tested with a tolerance of 0.005, 0.01 and 0.05.

#### Supplementary Figure 5

**Weakly contrasted and non-significantly different phenotypes between *Penicillium roqueforti* populations for various traits relevant for cheesemaking**

The color indicates assignment to the *Penicillium roqueforti* populations identified, as in the other figures. Horizontal lines on the boxplots represent the upper quartile, the median and the lower quartile. Dots represent the outlier values. Different letters indicate significant differences (Supplementary Table 4).**A** Growth on bread medium (colony size in mm after three days of growth) **B,C**: Spore production on cheese (B) and malt (C) media measured as optical density by spectrophotometer; **D**: Salt tolerance, i.e., growth rates on Petri dishes containing cheese and malt media with (8%) and without (0%) NaCl.

#### Supplementary Figure 6

**Enriched gene ontology (GO) terms for genes located within windows of 50kb showing both extreme divergence (1% upper quantile) between Roquefort and non-Roquefort populations and low diversity (5% lower quantile) within each of these two *Penicillium roqueforti* populations.**

Variation in color represents the -log10(p-value) of Fisher's exact test: the redder the more significant. The size of the box correlates with the number of occurrences of the GO term (minimum=3; maximum =42)**.**

#### Supplementary Table 1

**Description of the *Penicillium* strains analyzed in this study:** ID indicating the collection (LCP for laboratoire de cryptogamie, collection belonging to the National Museum of Natural History in Paris, LIP for laboratoire interprofessionnel de production d'Aurillac or UBOCC for université de Bretagne occidentale collection culture), species, environment and substrate of collection, assignation to the identified genetic clusters and accession numbers. In bold are strains with genome sequence available. Stars refer to genome assembly. The accession numbers correspond to the genome sequence or to the sequence of the two genetic markers used for strain genotyping and the references indicate the publications presenting these sequence data.

#### Supplementary Table 2

**Primer sets used in this study, with the marker name, its use, the analysis method, primer sequences and reference.** Their use was either to assign the *Penicillium roqueforti* strains without sequenced genome to the identified clusters, to determine if the genomic islands with presence/absence polymorphism detected in the sequenced genomes were present or absent in the strains, or to check the identity a specific strain in the cheese experiment. The amplicons were either sequenced or their presence/absence was assessed on agarose gels, as indicated in the analysis column.

#### Supplementary Table 3

**Phenotypic data for all strains.** Phenotypic data are given, for sporulation, proteolysis, lipolysis, and growth on Petri dishes. Missing data are marked with a cross (x) and unit of measurements are mentioned in between brackets. For growth measurement, speed is measured as the number of days it took for the molds to fill up a 3-cm diameter circle.

#### Supplementary Table 4

Results of statistical analyses performed for testing differences in nucleotidic diversity, genetic diversity, linkage disequilibrium and fitness traits between *Penicillium roqueforti* populations.

Models are detailed for each analysis/experiment, for the analyses of variance (ANOVA), Kruskal-Wallis rank tests, with response variables, explanatory variables (with their different classes in brackets) and their interactions when significant, Chi2 ($\chi 2$) values for Kruskal-Wallis tests or F-ratios for ANOVAs, degrees of freedom (df) and P values, with an asterisk and in bold when significant. Horizontal lines separate the different statistical models.

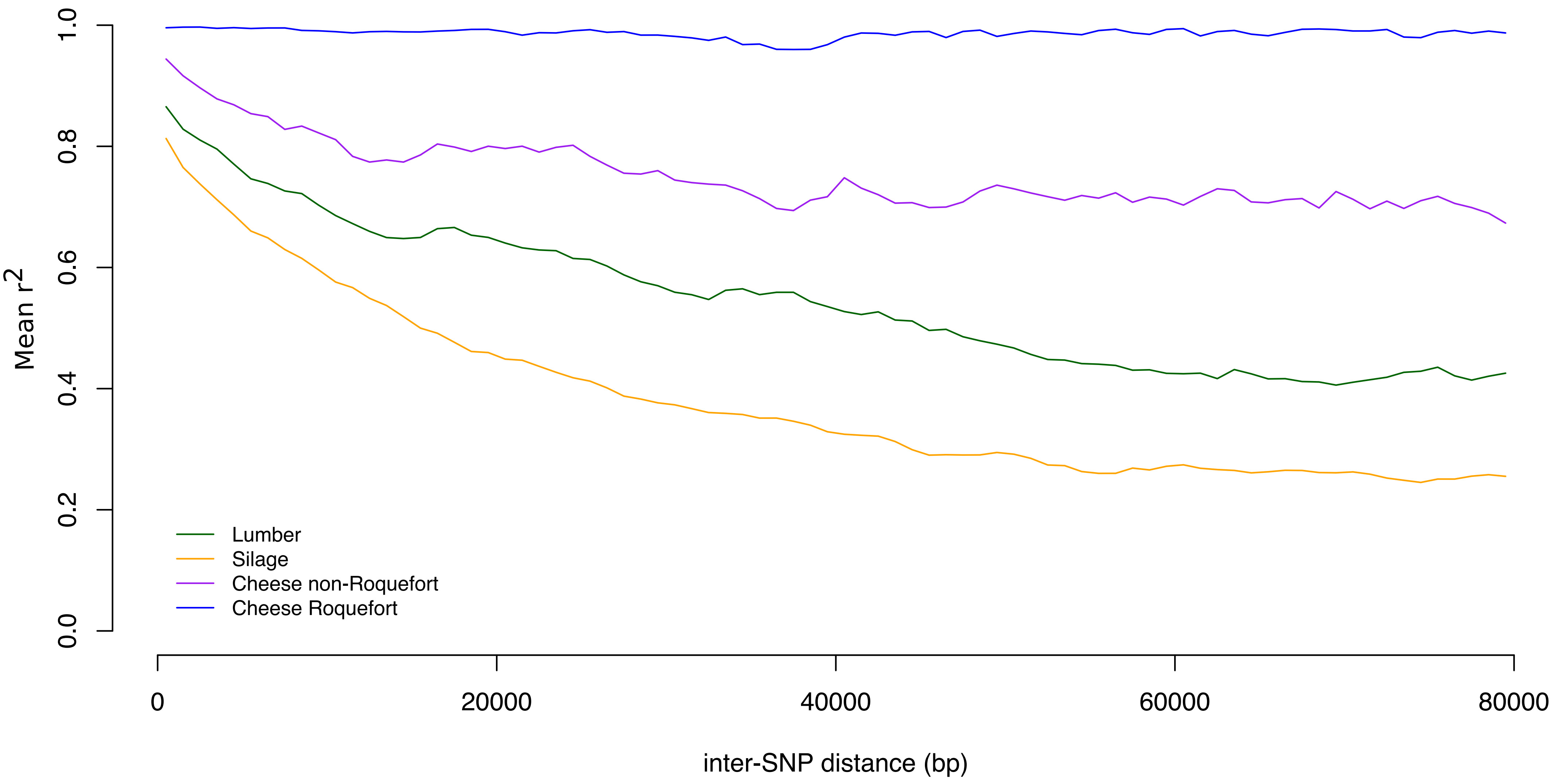

Mean r2
1.0
0.8
0.6
0.4
0.2
0.0
Lumber
Silage
Cheese non-Roquefort
Cheese Roquefort
0
20000
40000
60000
80000
inter-SNP distance (bp)

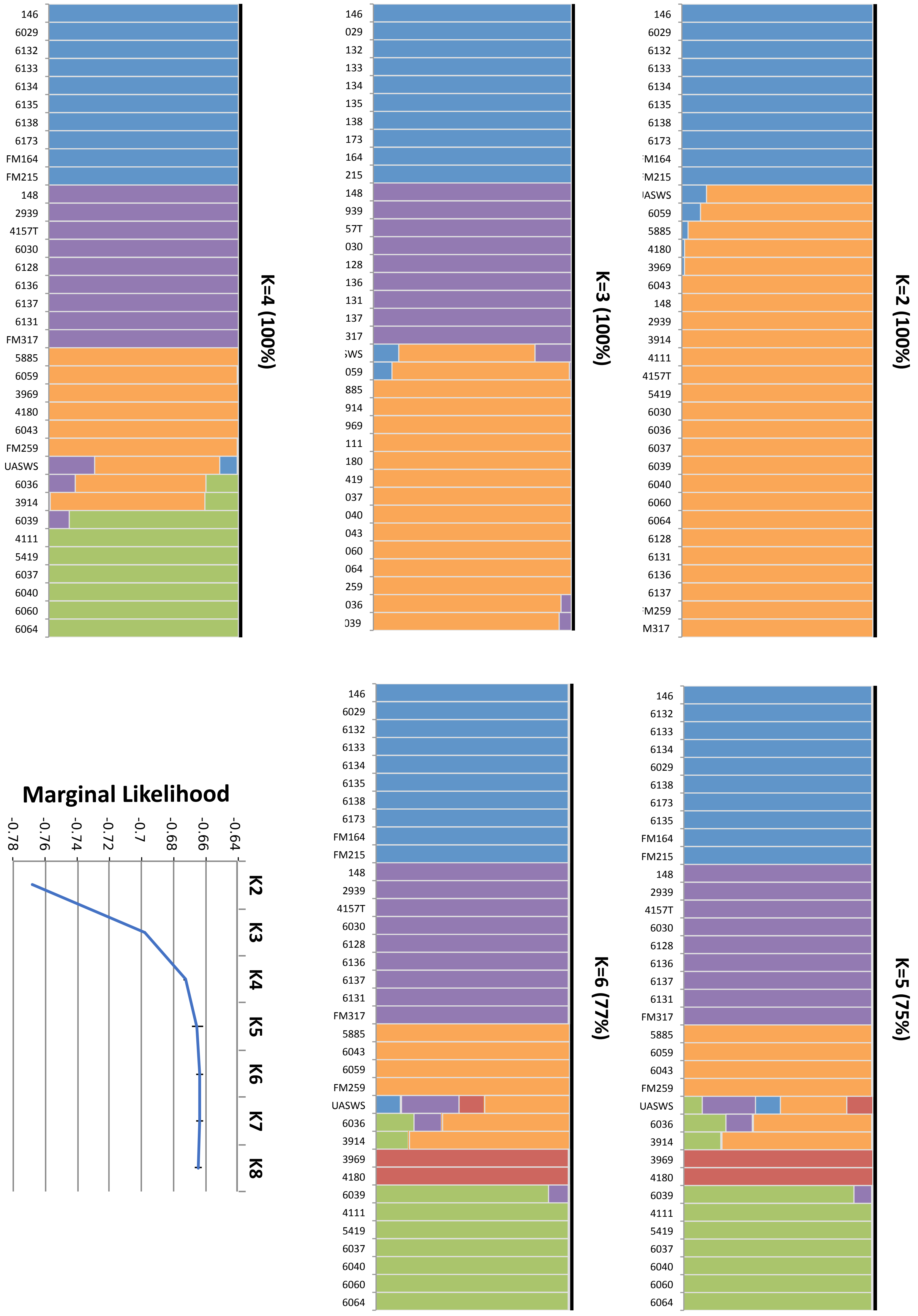

146
6029
6132
6133
6134
6135
6138
6173
FM164
FM215
148
2939
4157T
6030
6128
6136
6137
6131
FM317
5885
6059
3969
4180
6043
FM259
UASWS
6036
3914
6039
4111
5419
6037
6040
6060
6064
K=4 (100%)
146
029
132
133
134
135
138
173
164
215
148
939
57T
030
128
136
131
137
317
WS
059
885
914
969
111
180
419
037
040
043
060
064
259
036
039
K=3 (100%)
146
6029
6132
6133
6134
6135
6138
6173
M164
M215
ASWS
6059
5885
4180
3969
6043
148
2939
3914
4111
4157T
5419
6030
6036
6037
6039
6040
6060
6064
6128
6131
6136
6137
M259
M317
K=2 (100%)
Marginal Likelihood
-0.78
-0.76
-0.74
-0.72
-0.7
-0.68
-0.66
-0.64
K2
K3
K4
K5
K6
K7
K8
146
6029
6132
6133
6134
6135
6138
6173
FM164
FM215
148
2939
4157T
6030
6128
6136
6137
6131
FM317
5885
6043
6059
FM259
UASWS
6036
3914
3969
4180
6039
4111
5419
6037
6040
6060
6064
K=6 (77%)
146
6132
6133
6134
6029
6138
6173
6135
FM164
FM215
148
2939
4157T
6030
6128
6136
6137
6131
FM317
5885
6059
6043
FM259
UASWS
6036
3914
3969
4180
6039
4111
5419
6037
6040
6060
6064
K=5 (75%)

A

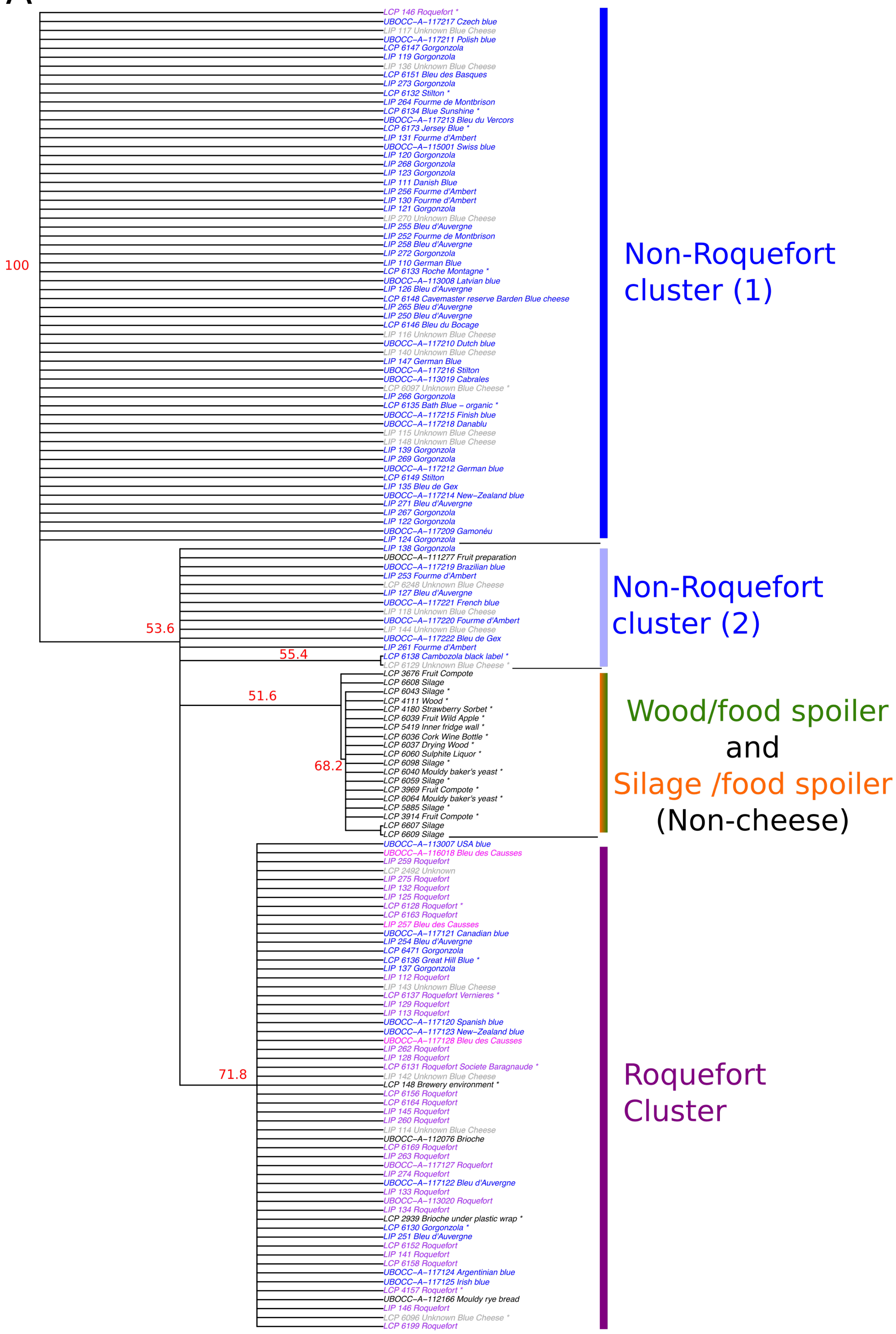


B

| Environment / Cluster | Roquefort and Bleu des Causses | All known cheeses except Roquefort and Bleu des Causses | Non-Cheese | Unknown blue cheeses |
|---|---|---|---|---|
| **Non-Roquefort (1)** | 1 | 51 | 0 | 8 |
| **Non-Roquefort (2)** | 0 | 9 | 1 | 4 |
| **Non-cheese** | 0 | 0 | 19 | 0 |
| **Roquefort** | 33 | 13 | 4 | 5 |

A

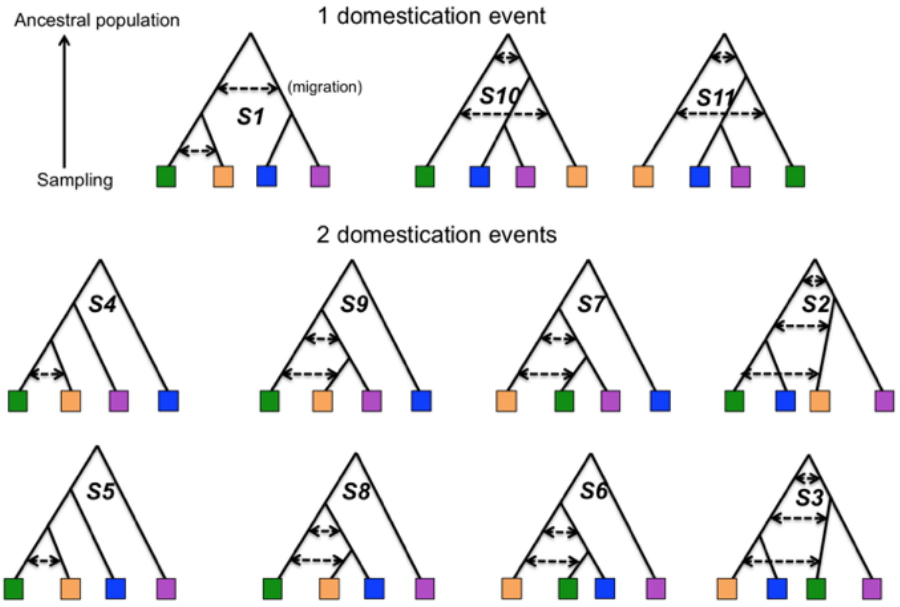


B

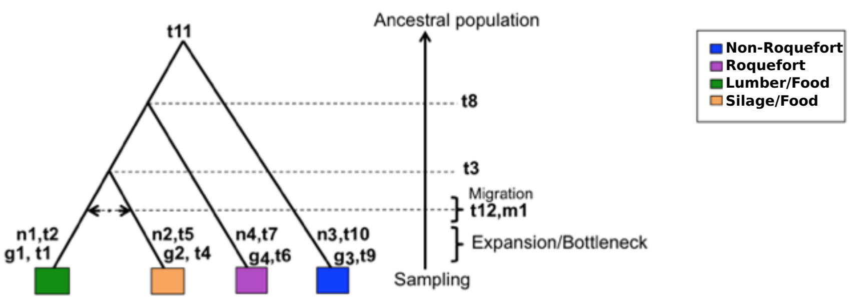


C

| Parameter | Description | Prior distribution |
|---|---|---|
| n1 | Bottleneck strength of lumber/food spoiler population | $10*(1\text{-}\log_{10}(unif(1,10^{0.9}))$ |
| n2 | Bottleneck strength of silage/food spoiler population | $10*(1\text{-}\log_{10}(unif(1,10^{0.9}))$ |
| n3 | Bottleneck strength of non-Roquefort population | $10*(1\text{-}\log_{10}(unif(1,10^{0.9}))$ |
| n4 | Bottleneck strength of Roquefort population | $10*(1\text{-}\log_{10}(unif(1,10^{0.9}))$ |
| g1 | Growth rate of lumber/food spoiler population after bottleneck | unif(0,10000) |
| g2 | Growth rate of silage/food spoiler population after bottleneck | unif(0,10000) |
| g3 | Growth rate of non-Roquefort population after bottleneck | unif(0,10000) |
| g4 | Growth rate of Roquefort population after bottleneck | unif(0,10000) |
| m | Migration rate between lumber/food spoiler and silage/food spoiler populations before bottleneck | $1\text{-}\log_{1000}(unif(\sqrt{1000},1000))$ |
| t1 | Time backward since expansion of lumber/food spoiler population | $1\text{-}\log_{10}(unif(10^{1\text{-}t2},10))$ |
| t2 | Time backward since bottleneck of lumber/food spoiler population | $1\text{-}\log_{10}(unif(10^{1\text{-}t12},10))$ |
| t3 | Time backward to population split between lumber/food spoiler and silage/food spoiler ancestral populations | $1\text{-}\log_{10}(unif(10^{1\text{-}t8},10))$ |
| t4 | Time backward since expansion of lumber/food spoiler population | $1\text{-}\log_{10}(unif(10^{1\text{-}t5},10))$ |
| t5 | Time backward since bottleneck of lumber/food spoiler population | $1\text{-}\log_{10}(unif(10^{1\text{-}t12},10))$ |
| t6 | Time backward since expansion of non-Roquefort population | $1\text{-}\log_{10}(unif(10^{1\text{-}t7},10))$ |
| t7 | Time backward since bottleneck of non-Roquefort population | $1\text{-}\log_{10}(unif(10^{1\text{-}t8},10))$ |
| t8 | Time backward to population split between non-cheese ancestral population and Roquefort ancestral population | $1\text{-}\log_{10}(unif(10^{1\text{-}t11},10))$ |
| t9 | Time backward since expansion of Roquefort population | $1\text{-}\log_{10}(unif(10^{1\text{-}t10},10))$ |
| t10 | Time backward since bottleneck of Roquefort population | $1\text{-}\log_{10}(unif(10^{1\text{-}t11},10))$ |
| t11 | Time backward to population split between non-cheese and Roquefort ancestral population and non-Roquefort ancestral population | $1\text{-}\log_{10}(unif(1,10))$ |
| t12 | Time backward when migration between lumber/food spoiler and silage/food spoiler populations occurred | $1\text{-}\log_{10}(unif(10^{1\text{-}t3},10))$ |

D

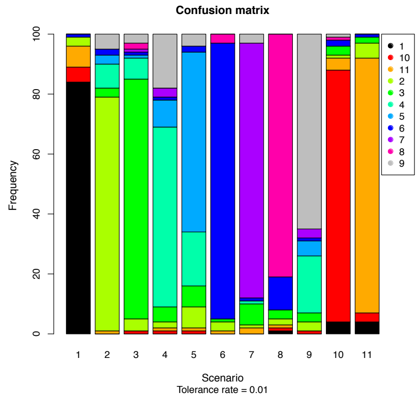


E

| Parameter | Mode | Mean | Median | Credibility intervals |
|---|---|---|---|---|
| **n1** | 2.0149 | 4.11 | 3.5888 | (1.1021 - 9.4243) |
| **n2** | 1.5876 | 3.0058 | 2.2995 | (1.0365 - 8.7107) |
| **n3** | 1.8624 | 3.9409 | 3.3063 | (1.0827 - 9.1958) |
| **n4** | 1.8263 | 3.9577 | 3.3681 | (1.0849 - 9.3731) |
| **g1** | 2820.5988 | 4961.3045 | 4986.4674 | (231.8938 - 9691.0923) |
| **g2** | 3905.3683 | 4718.1456 | 4491.3869 | (188.2575 - 9725.8372) |
| **g3** | 2909.4405 | 4993.1661 | 4852.604 | (336.8148 - 9677.2838) |
| **g4** | 780.5916 | 3718.1832 | 3169.6466 | (95.9802 - 9498.3444) |
| **m** | 0.0406 | 0.1244 | 0.0916 | (0.0032 - 0.4089) |
| **t1** | 0.0001 | 0.004 | 0.0007 | (0 - 0.0267) |
| **t2** | 0.0005 | 0.0082 | 0.0023 | (0 - 0.0478) |
| **t3** | 0.0068 | 0.038 | 0.0163 | (0 - 0.2099) |
| **t4** | 0.0002 | 0.0039 | 0.0009 | (0 - 0.027) |
| **t5** | 0.0006 | 0.0079 | 0.0023 | (0 - 0.0454) |
| **t6** | 0.001 | 0.0061 | 0.0019 | (0.0005 - 0.04) |
| **t7** | 0.0045 | 0.0249 | 0.0097 | (0.0012 - 0.1402) |
| **t8** | 0.0181 | 0.0816 | 0.0419 | (0.0023 - 0.3985) |
| **t9** | 0.0096 | 0.0523 | 0.0242 | (0.0017 - 0.268) |
| **t10** | 0.0268 | 0.1173 | 0.0662 | (0.0045 - 0.4939) |
| **t11** | 0.0772 | 0.2675 | 0.1875 | (0.017 - 0.8711) |
| **t12** | 0.0019 | 0.0169 | 0.0062 | (0 - 0.0865) |

| Scenario | Rejection | | | | Logistic Regression | | | | Neural Network | | | |
|---|---|---|---|---|---|---|---|---|---|---|---|---|
| | **Tol= 0.005** | **Tol= 0.01** | **Tol= 0.05** | **Tol= 0.1** | **Tol= 0.005** | **Tol= 0.01** | **Tol= 0.05** | **Tol= 0.1** | **Tol= 0.005** | **Tol= 0.01** | **Tol= 0.05** | **Tol= 0.1** |
| 1 | 0.0817 | 0.0528 | 0.0133 | 0.0076 | 0 | 0 | 0 | 0 | 0.0091 | 0 | 0 | 0 |
| 2 | 0.0781 | 0.0504 | 0.0506 | 0.0821 | 0 | 0 | 0.0006 | 0.0001 | 0.0297 | 0.0181 | 0 | 0 |
| 3 | 0.0723 | 0.0471 | 0.0491 | 0.0824 | 0 | 0 | 0 | 0 | 0.0301 | 0.0168 | 0 | 0 |
| **4** | **0.1294** | **0.2017** | **0.3283** | **0.3086** | 0.002 | 0.0036 | 0.1199 | 0.0028 | 0.2394 | **0.4518** | 0.464 | **0.5601** |
| **5** | 0.0737 | 0.0638 | 0.1411 | 0.1641 | **0.9889** | 0.0427 | **0.5678** | **0.9965** | **0.5288** | 0.4141 | **0.5356** | 0.4118 |
| 6 | 0.0818 | 0.0522 | 0.012 | 0.0064 | 0 | 0.0002 | 0.2255 | 0 | 0.009 | 0 | 0 | 0 |
| 7 | 0.0966 | 0.1281 | 0.0615 | 0.0431 | 0.009 | **0.9534** | 0.084 | 0.0006 | 0.0078 | 0.0003 | 0 | 0 |
| 8 | 0.0956 | 0.0973 | 0.0386 | 0.0251 | 0 | 0 | 0.0022 | 0 | 0.0089 | 0 | 0 | 0 |
| 9 | 0.123 | 0.1994 | 0.2806 | 0.2676 | 0 | 0 | 0 | 0 | 0.1181 | 0.099 | 0.0003 | 0.028 |
| 10 | 0.0821 | 0.0529 | 0.0122 | 0.0065 | 0 | 0 | 0 | 0 | 0.0093 | 0 | 0 | 0 |
| 11 | 0.0858 | 0.0544 | 0.0125 | 0.0066 | 0 | 0 | 0 | 0 | 0.0097 | 0 | 0 | 0 |

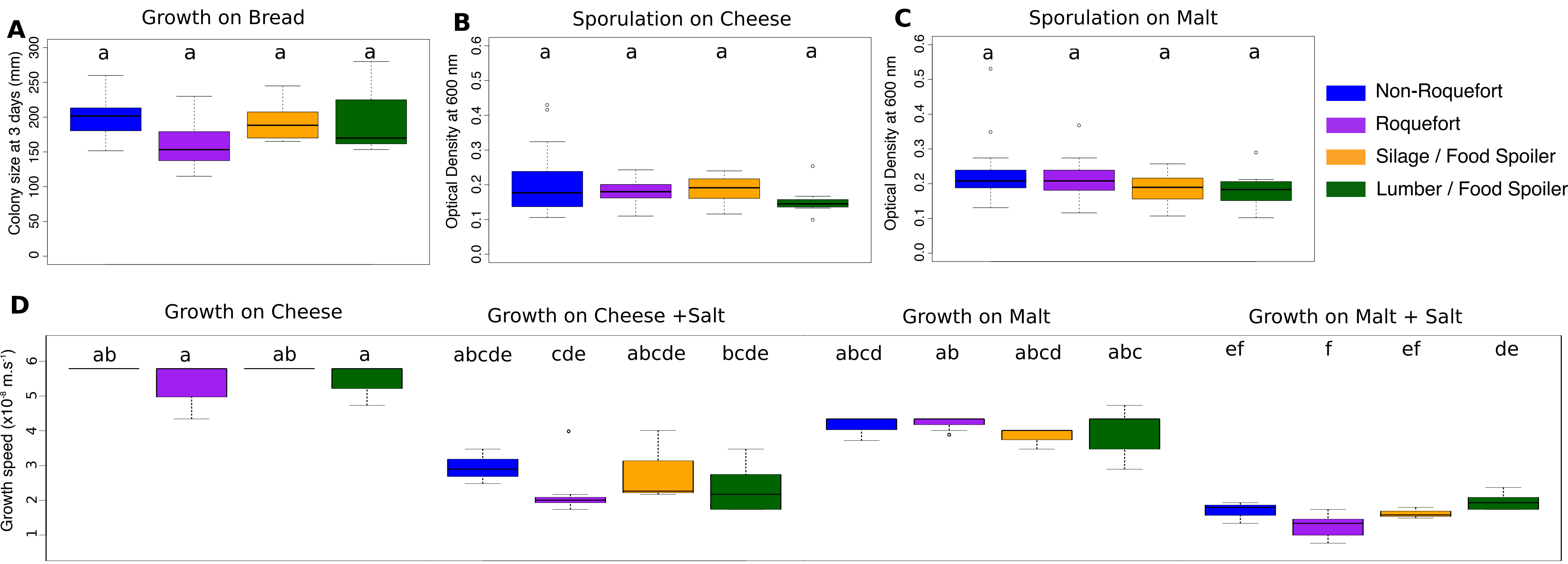
A
Growth on Bread
Colony size at 3 days (mm)
0
50
100
150
200
250
300
a
a
a
a
B
Sporulation on Cheese
Optical Density at 600 nm
0.0
0.1
0.2
0.3
0.4
0.5
0.6
a
a
a
a
C
Sporulation on Malt
Optical Density at 600 nm
0.0
0.1
0.2
0.3
0.4
0.5
0.6
a
a
a
a
Non-Roquefort
Roquefort
Silage / Food Spoiler
Lumber / Food Spoiler
D
Growth on Cheese
Growth on Cheese +Salt
Growth on Malt
Growth on Malt + Salt
Growth speed (x10-8 m.s-1)
1
2
3
4
5
6
ab
a
ab
a
abcde
cde
abcde
bcde
abcd
ab
abcd
abc
ef
f
ef
de

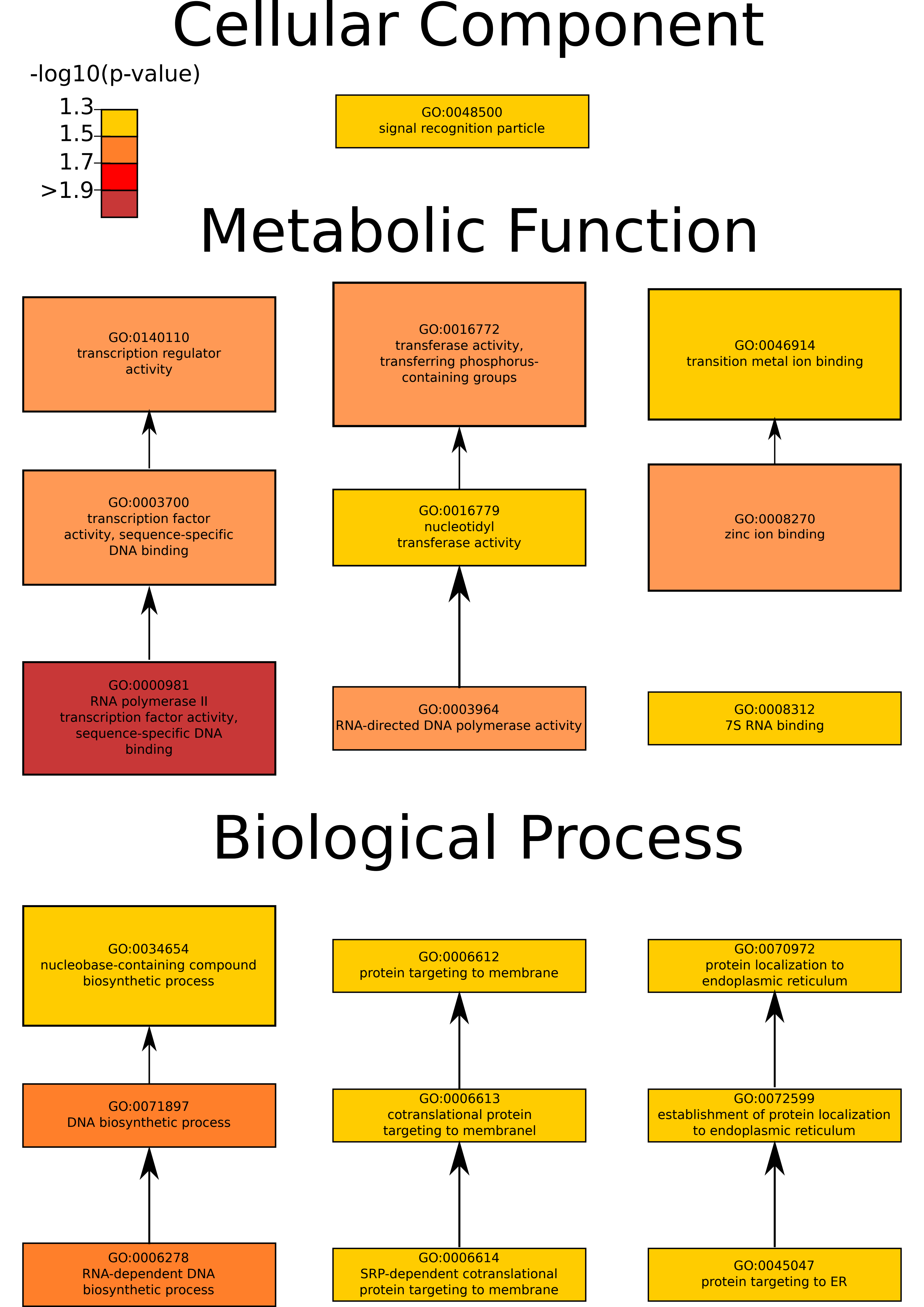

Cellular Component
-log10(p-value)
1.3
1.5
1.7
>1.9
GO:0048500
signal recognition particle
Metabolic Function
GO:0140110
transcription regulator activity
GO:0016772
transferase activity, transferring phosphorus-containing groups
GO:0046914
transition metal ion binding
GO:0003700
transcription factor activity, sequence-specific DNA binding
GO:0016779
nucleotidyl transferase activity
GO:0008270
zinc ion binding
GO:0000981
RNA polymerase II transcription factor activity, sequence-specific DNA binding
GO:0003964
RNA-directed DNA polymerase activity
GO:0008312
7S RNA binding
Biological Process
GO:0034654
nucleobase-containing compound biosynthetic process
GO:0006612
protein targeting to membrane
GO:0070972
protein localization to endoplasmic reticulum
GO:0071897
DNA biosynthetic process
GO:0006613
cotranslational protein targeting to membranel
GO:0072599
establishment of protein localization to endoplasmic reticulum
GO:0006278
RNA-dependent DNA biosynthetic process
GO:0006614
SRP-dependent cotranslational protein targeting to membrane
GO:0045047
protein targeting to ER
